# Editing Efficiency Across Crop Families: A Systematic Review and Meta-Analysis of CRISPR/SpCas9 Knockout Outcomes in Cucurbitaceae, Brassicaceae, Solanaceae and Poaceae

**DOI:** 10.64898/2026.07.03.736175

**Authors:** Yemisi O. Olagunju, Mobolaji T. Oladunjoye

## Abstract

Reported CRISPR/SpCas9 editing efficiencies in crops span 0–100%, but no quantitative synthesis has separated taxonomic family from delivery method, ploidy, clustering or publication bias. This meta-analysis estimated pooled per-T0-line editing efficiency across Cucurbitaceae, Brassicaceae, Solanaceae and Poaceae, and tested whether family is an independent moderator after adjustment for delivery and ploidy. A PRISMA 2020 systematic review identified peer-reviewed studies using SpCas9 with extractable per-line T0 edit counts; data were extracted independently by two reviewers, with inter-rater agreement reported. Logit proportions were synthesised with a binomial-normal generalised linear mixed model, and the family-as-moderator hypothesis was tested by a small-sample CR2 cluster-robust F-test on a three-level model with study-level clustering. Publication bias was assessed by Egger’s regression and trim-and-fill. Twenty-two studies contributed 172 per-line effect sizes (Cucurbitaceae k=14, Brassicaceae k=20, Solanaceae k=68, Poaceae k=70). Pooled editing efficiency was 61.8% (95% CI 54.5–68.6%) with I²=93.4% (τ²=3.21) and a 95% prediction interval of approximately 5–98%. Per-family estimates ranged from 47.8% (Poaceae) to 73.8% (Brassicaceae); the univariate Q test was significant (p=0.0016), but family did not survive cluster-robust adjustment (F=0.73, p=0.63). Intraclass correlation placed 64.4% of variance at the study level, and Solanaceae remained dominated by a single study (58/68 rows). Funnel asymmetry was severe (Egger p<0.0001), and trim-and-fill reduced the bias-adjusted estimate to 45.2% (95% CI 39.0–51.5%). Apparent crop-family differences dissolve once within-study clustering and methodological covariates are accounted for; the bias-adjusted pooled estimate is closer to 45% than to 62%, and reported editing efficiencies reflect study-level factors more than taxonomic family.

**Key Message:** Apparent between-family differences in CRISPR/SpCas9 editing efficiency across four crop families reflect within-study clustering and publication bias, not intrinsic biology; family is not an independent moderator after cluster-robust adjustment.

## 1. Introduction

### 1.1 The editing-efficiency problem

CRISPR/SpCas9 is now a default tool for targeted mutagenesis in crops, but the proportion of regenerated lines that carry a confirmed targeted edit varies enormously among published reports. Within the same nuclease and target type, edit-positive proportions in T0 lines have been reported from below 10% to 100% across cucurbits, brassicas, solanaceous crops and grasses (Feng et al. 2021; Hooghvorst et al. 2019; Howells et al. 2018; Pan et al. 2022; Tian et al. 2017; Wang et al. 2024; Xin et al. 2022). The published range is wide enough that no single number captures “typical” efficiency, and practical questions, how many transformants to attempt, how aggressively to screen, when to switch delivery system, are answered ad hoc per laboratory.

Two broad classes of explanation for this dispersion compete in the literature. The first attributes variation primarily to taxonomic and genome-level features: ploidy, genome size, transformation amenability, gene redundancy through paralogy, and culture-system regeneration efficiency (Ma et al. 2019; Yang et al. 2017; Yang et al. 2018; Zheng et al. 2020). The second attributes the variation to method-level features that are confounded with taxonomy: delivery vehicle (Agrobacterium versus protoplast PEG versus biolistic), sgRNA-promoter architecture, detection assay sensitivity, and decisions about which lines enter the screened denominator (Brooks et al. 2014; Gonzalez et al. 2021; Pan et al. 2016; Zhang et al. 2020). The two explanations make different operational predictions, a family signal that survives adjustment for method versus a family signal that vanishes after adjustment, but they have not been formally separated in a synthesis that uses all four families at once.

### 1.2 Why a quantitative cross-family synthesis is needed now

Family-specific reviews exist for cucurbits (Feng et al. 2023), cereals (Char et al. 2017; Lee et al. 2019) and Solanaceae (Marik et al. 2024), and single laboratories have compared two or three species from a single family under controlled conditions (Gasparis et al. 2018; Lawrenson et al. 2024; Milner et al. 2024). None of these has pooled effect sizes across families, formally tested the family-versus-method confound, or quantified publication bias on the proportion scale using the binomial-normal generalised linear mixed model that is now standard for sparse-event proportion meta-analysis (Stijnen et al. 2010; Viechtbauer 2010). The methodological vocabulary used to handle within-study dependence, multilevel random-effects models with cluster-robust variance, has matured but is not yet routine in plant-biotechnology synthesis (Pustejovsky and Tipton 2018).

The cost of the gap is not theoretical. Laboratories starting a new species, or moving an established construct into a new family, currently extrapolate from one published study at a time. Reviewers and funders evaluating CRISPR-based breeding proposals lack a defensible “typical efficiency, with uncertainty” figure to anchor expectations. The wider field is also vulnerable to publication-bias dynamics that are particularly acute in transformation work, where laboratories rarely publish constructs that failed to produce edits at all and small studies are disproportionately likely to appear when efficiency is high (Sterne et al. 2011).

### 1.3 Objectives and hypothesis

This synthesis was designed to address three pre-specified objectives:

**O1.** Estimate the pooled proportion of regenerated/T0 lines carrying a confirmed targeted SpCas9 edit, overall and within each of Cucurbitaceae, Brassicaceae, Solanaceae and Poaceae, using a binomial-normal GLMM on raw counts.

**O2.** Quantify between-family heterogeneity and test whether crop family is an independent moderator after adjustment for delivery method and ploidy (hypothesis H1).

**O3.** Identify the residual sources of variation, within-study clustering, publication bias, and influential single studies, most likely to mislead future readers of the literature.

**Hypothesis H1.** Crop family explains residual editing-efficiency heterogeneity beyond what is captured by delivery method and ploidy. The pre-specified test is a small-sample cluster-robust F-test (HTZ) on a three-level rma.mv fit, with a nested likelihood-ratio statistic on the corresponding GLMM as supporting evidence.

## 2. Methods

### 2.1 Protocol, registration and reporting

The protocol was prepared in PRISMA-P format and registered on the Open Science Framework (osf.io/vgf5u). Reporting follows the PRISMA 2020 statement (Page et al. 2021); the completed PRISMA 2020 checklist accompanies this manuscript as Supplementary File S1. The analysis dataset (Data_Extraction_v4_VERIFIED.csv), the structured extraction workbook (v4 dual-reviewer-verified), the R analysis script (CRISPR_meta_analysis.R), and the second-reviewer verification report (Supplementary File S8) are deposited at (https://doi.org/10.17605/OSF.IO/7T3SF); results are exactly reproducible from these four files under set.seed(20250101) on R ≥ 4.5.1 with metafor 4.x and clubSandwich 0.5.x.

### 2.2 Eligibility criteria

Eligible studies were peer-reviewed primary research articles reporting CRISPR/SpCas9 targeted mutagenesis in at least one species belonging to Cucurbitaceae, Brassicaceae, Solanaceae or Poaceae, with extractable counts of T0 (regenerated) lines confirmed to carry an edit at the targeted locus relative to the total number of T0 lines screened by a molecular detection assay. Studies using nucleases other than SpCas9 (e.g. LbCas12a, base editors, prime editors) were excluded, as were protoplast-only transient-edit-rate studies in which no plants were regenerated, narrative or systematic reviews without primary effect sizes, and abstracts/preprints not subsequently peer-reviewed. T1-generation rates, per-locus rates and per-allele rates, where reported in lieu of per-T0-line rates, were extracted but flagged as primary_outcome_flag = “No” and reserved for sensitivity analysis only.

### 2.3 Information sources and search

Searches were executed in PubMed/MEDLINE, Web of Science Core Collection, Scopus, Google Scholar (first 200 records), bioRxiv, CAB Abstracts and AGRIS. The full Boolean strings and database-specific MeSH/Emtree adaptations are listed in Supplementary File S2; final search date 23 June 2026. Forward and backward snowballing was performed on each included study. Figure 1 shows the PRISMA 2020 study-selection process. The search returned 3,710 records across the seven databases, plus 12 additional records identified through forward and backward snowballing, giving 3,722 records in total (per-database hit counts and full Boolean search strings are provided in Supplementary File S2). Deduplication in Zotero removed 1,570 records (∼42%), a rate consistent with expected overlap between AGRIS and Web of Science/CAB Abstracts, both of which extensively index agricultural plant-science literature. The remaining 2,152 unique records were screened at title/abstract level, of which 2,089 were excluded and 63 progressed to full-text assessment. Forty studies were excluded at full-text stage for the reasons shown in Figure 1, leaving 23 studies for qualitative synthesis. Of these, 21 were dual-reviewer-verified as of protocol version 4 (v4); Lawrenson (2024) and Milner (2024) were added post hoc as version 5 (v5) following identification through snowballing. Twenty-two studies contributed primary per-T0-line effect sizes (k = 172) to the meta-analytic model. The remaining study, Ma X (2015), reports effect sizes per locus rather than per T0 line and was therefore restricted to the sensitivity analysis (4 effect sizes); combined with 3 T1-generation rows from Zhang Z (2019), this yielded 7 sensitivity-only effect sizes evaluated separately from the primary dataset.

**Figure 1.**
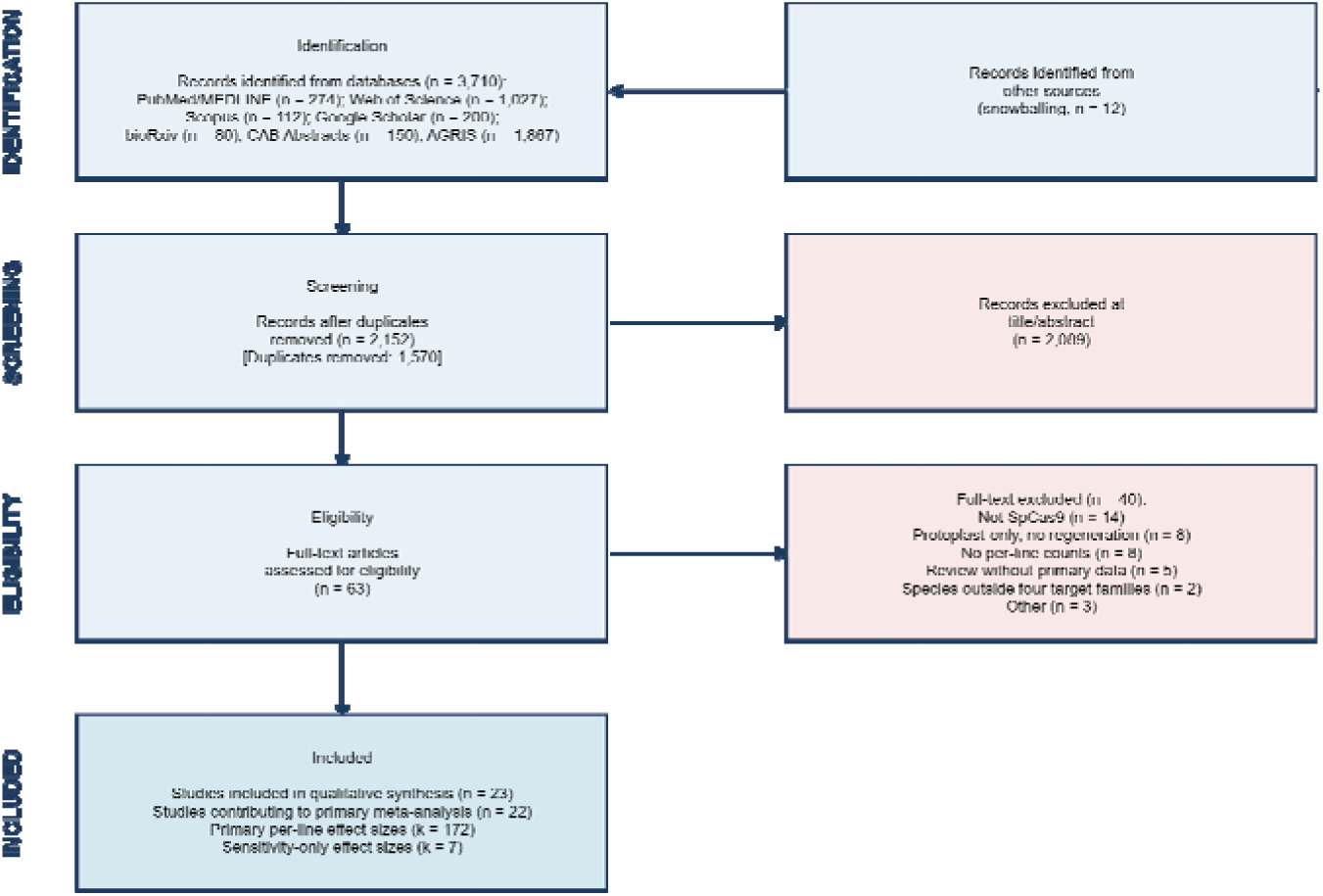
PRISMA 2020 study-selection flow diagram. Searches of seven databases plus citation snowballing (23 June 2026) identified 3,722 records. After deduplication (n = 1,570) and title/abstract screening, 63 studies underwent full-text assessment; 23 met inclusion criteria, of which 22 contributed effect sizes (k = 172) to the primary meta-analytic model. Supplementary File S2 lists per-database hit counts and search strings.

### 2.4 Study selection and data extraction (dual-reviewer)

Title and abstract screening was performed independently by two reviewers (Y.O. Olagunju, NIHORT, Ibadan; and M.T. Oladunjoye, NIHORT, Ibadan); inter-rater agreement at this stage was perfect (Cohen’ κ = 1.000, N = 21 studies). Full-text data extraction was similarly performed independently by both reviewers using the structured workbook described in Supplementary File S5. Inter-rater agreement was near-perfect across all extracted variables: κ = 1.000 for ploidy, delivery method, and detection assay; κ = 0.953 for primary-outcome flag and generation coding; and 100% exact agreement on numerator and denominator counts (N = 129 rows). The sole substantive disagreement concerned rows E0131–E0133 (Zhang Z 2019), which the first reviewer retained in primary synthesis with generation coded as a moderator, and the second reviewer reclassified as sensitivity-only on the grounds that T1-generation data are not interchangeable with the per-T0-line primary outcome definition. Following structured reconciliation, these three rows were moved to sensitivity analysis, reducing the primary synthesis dataset from k = 129 to k = 126 effect sizes. Detailed kappa tabulation by variable and full reconciliation minutes are in the verification report (Supplementary File S8).

Extracted variables comprised species, cultivar, ploidy, gene/locus, construct details (Cas promoter, sgRNA promoter, sgRNA processing architecture, booster), transformation method, explant tissue, selection marker, target class, numerator (T0 lines confirmed edited), denominator (T0 lines screened), generation, biallelic counts where reported, detection assay, primary-outcome flag, per-line vs per-locus distinction, and a missing-data route. Six rows have numerators or denominators reconstructed from reported percentages multiplied by N where exact counts were not tabulated (Zheng 2020 E0101–E0102 from per-allele rates; Zhang Z 2019 E0131–E0133 from efficiency-percentage × N; Yang Y 2018 E0098 where two text values differed by 0.1 percentage point). These reconstructions were verified independently by both reviewers; E0131–E0133 were subsequently reclassified to sensitivity-only as noted above.

### 2.5 Outcome definition and harmonisation

The primary outcome is the per-T0-line edit proportion: numerator = the number of regenerated T0 lines confirmed by sequencing or sequencing-equivalent assay to carry an indel at the target locus; denominator = the total number of T0 lines screened. When studies reported multiple constructs targeting the same gene in the same genotype with the same delivery system, counts were pooled into one row; otherwise one row was extracted per construct. Ma X 2015 reports per-locus rates only and was excluded from primary synthesis; the per-line restriction is sensitivity-tested by re-fitting the primary GLMM with 3Ma X 2015 per-locus rows added (Section 3.5). The Zhang Z 2019 T1-generation rows are reserved for the same sensitivity test.

### 2.6 Statistical analysis

The primary model was a binomial-normal generalised linear mixed model implemented through metafor::rma.glmm with measure = “PLO” (logit-transformed proportion) and method = “ML” with adaptive Gauss–Hermite quadrature (nAGQ = 7), following the Stijnen et al. (2010) parameterisation that operates on raw n and N counts and does not require continuity correction. Pooled proportions were obtained on the logit scale and back-transformed for reporting. Heterogeneity was summarised by τ², I², the residual Q test, and the 95% prediction interval (IntHout et al. 2016). The per-family model was fitted within each family with the same specification.

Because many rows came from the same study (Zhang N 2019 alone contributes 58 of 68 Solanaceae rows), a three-level multilevel meta-analytic model was fitted using metafor::rma.mv with random structure ∼ 1 | study_id / row_id, after first transforming counts to logit proportions through escalc(measure = “PLO”, add = 0.5, to = “only0”) so that boundary cases received continuity correction without affecting interior cases. CR2 cluster-robust standard errors and small-sample Wald (HTZ) tests were obtained via clubSandwich::vcovCR and Wald_test, clustering by study_id (Pustejovsky and Tipton 2018). This multilevel rma.mv fit is the basis of the hypothesis test for H1: model M1 included delivery and ploidy as covariates, and model M2 added family; the cluster-robust HTZ F-test on the joint nullity of the three family contrasts in M2 is the registered test of H1.

Subgroup analyses, fitted as moderator additions to the GLMM, were pre-specified for family, harmonised delivery method, harmonised ploidy, target class, sgRNA-promoter system, detection assay and generation. Heterogeneous text codings of moderators in the extraction workbook were collapsed to analysable factors using the rules in Supplementary Table S4; the original verbatim codings were preserved.

### 2.7 Sensitivity, influence and publication bias

Sensitivity analyses comprised: (i) leave-one-study-out on the overall pooled estimate; (ii) re-fitting the Solanaceae estimate after excluding Zhang N 2019; (iii) extension of the primary dataset to include the 7 sensitivity-only rows (Ma X 2015 per-locus rows and Zhang Z 2019 T1 wheat rows). Influence diagnostics (Cook’s distance, hat values, the Baujat plot) were generated from a univariate REML rma fit on the escalc-transformed data. Publication-bias diagnostics were Egger’s regression of standardised effect on its standard error (Egger et al. 1997) and the Duval and Tweedie trim-and-fill estimator (Duval and Tweedie 2000); both are interpreted with the field-specific caveat that single-laboratory transformation studies with low or zero edit rates are routinely unpublished.

## 3. Results

### 3.1 Studies included and dataset structure

Twenty-two primary-research studies provided 172 per-line effect sizes that met the eligibility criteria and entered the primary synthesis (Table 1; full study-level details in Supplementary Table S5). Twenty of these studies were dual-reviewer-verified (κ = 0.953–1.000 across extracted variables) at v4. The two further open-access studies (Lawrenson 2024, 16 rows from barley and wheat; Milner 2024, 30 rows from rice, barley and wheat) were extracted from full-text post hoc and added to the workbook as v5; second-reviewer verification of these 46 new rows is pending. A further 7 effect sizes (T1-generation rows from Zhang Z 2019; per-locus rows from Ma X 2015) are retained as sensitivity-only and excluded from primary pooling per the pre-registered protocol.

Cucurbitaceae contributed 14 effect sizes from 6 studies (Tian 2017; Hooghvorst 2019; Pan 2022; Wang 2024; Xin 2022; Feng 2021); Brassicaceae 20 effect sizes from 4 studies (Yang H 2017; Ma C 2019; Yang Y 2018; Zheng 2020); Solanaceae 68 effect sizes from 4 studies (Brooks 2014; Pan 2016; Zhang N 2019; Gonzalez 2021), of which 58 came from Zhang N 2019’s disease-resistance gene screen; and Poaceae 70 primary effect sizes from 8 studies (Howells 2018; Gasparis 2018; Zhang H 2014; Char 2017; Lee 2019; Zhang Z 2019; Lawrenson 2024; Milner 2024), with three additional T1-generation Zhang Z 2019 rows and four Ma X 2015 per-locus rows reserved for sensitivity. The dataset spans diploid, autotetraploid, allotetraploid and hexaploid genomes; Agrobacterium-mediated delivery dominated (k = 170) with a small protoplast PEG/RNP arm from Gonzalez 2021 (k = 2).

### 3.2 Overall pooled editing efficiency

The binomial-normal GLMM on the full primary dataset (k = 172 effect sizes, 22 studies) estimated a pooled SpCas9 editing efficiency of **61.8% (95% CI 54.5–68.6%)** at the per-T0-line level. Residual heterogeneity was severe (I² = 93.4%, τ² = 3.21 on the logit scale) and the 95% prediction interval, which describes where a new study’s pooled estimate is expected to fall conditional on the same effects governing the included literature, spanned approximately 4.6–98.2%, effectively the entire unit interval. In operational terms, the pooled point estimate by itself provides essentially no information for any specific upcoming transformation experiment; the prediction interval captures the meaningful uncertainty. Heterogeneity has, if anything, increased relative to the v4 dataset (I² = 89.8%, τ² = 1.99 at k = 126), driven by the Milner 2024 entries that span 0% to 60% within a single gene target across three Poaceae species and two guide architectures.

The three-level rma.mv fit (random structure: row in study) returned a pooled estimate of 56.6% (95% CI 43.0–69.3%) and partitioned the residual heterogeneity sharply: σ²(study-level) = 1.309, σ²(row-level) = 0.724, giving an intraclass correlation of 64.4%. Almost two-thirds of the variance in logit edit proportions sits between studies rather than between effect sizes within a study, a numerical fingerprint of strong laboratory and construct-batch effects. The ICC has fallen slightly from 73.3% at v4, reflecting the addition of 46 rows from two new laboratories (Lawrenson at JIC, Milner at NIAB) that contribute new between-study variance, but the qualitative picture is unchanged. The implication for the H1 test in Section 3.4 is that any analysis that treats the 172 rows as independent observations will substantially underestimate the standard errors of moderator coefficients.

### 3.3 Per-family pooled effects

Per-family binomial-normal GLMM fits (Table 2; forest plot Figure 2) returned pooled estimates ranging from 47.8% in Poaceae to 73.8% in Brassicaceae. Solanaceae sat at 71.1% with the narrowest confidence interval of any family (63.0–78.0%), reflecting the weight of Zhang N 2019’s 58 effect sizes; Cucurbitaceae sat at 52.0% (30.3–73.0%) with the widest interval of any well-populated family. The 26-percentage-point spread between the lowest and highest pooled family estimates is visually striking. Of note, the Cucurbitaceae, Brassicaceae and Solanaceae pooled estimates and confidence intervals are unchanged from v4 because the underlying rows are identical; only Poaceae has shifted, from 52.6% on 24 rows from 6 studies (v4) to 47.8% on 70 rows from 8 studies (v5), as the Lawrenson and Milner entries together pull the Poaceae arm downward despite Lawrenson contributing high-efficiency wheat and barley data. The Poaceae arm now also exhibits the highest within-family heterogeneity of the four families (I² = 95.9%, τ² = 4.25), reflecting genuine biological dispersion between Milner’s near-zero ribozyme entries and Lawrenson’s near-100% architecture-A barley entries within the same crop family. Brassicaceae’s within-family heterogeneity remains in large part driven by the Yang Y 2018 construct-by-construct screen of CLAVATA genes in oilseed rape, where only 5 of 10 sgRNAs produced any edits in T0 plants.

**Figure 2.**
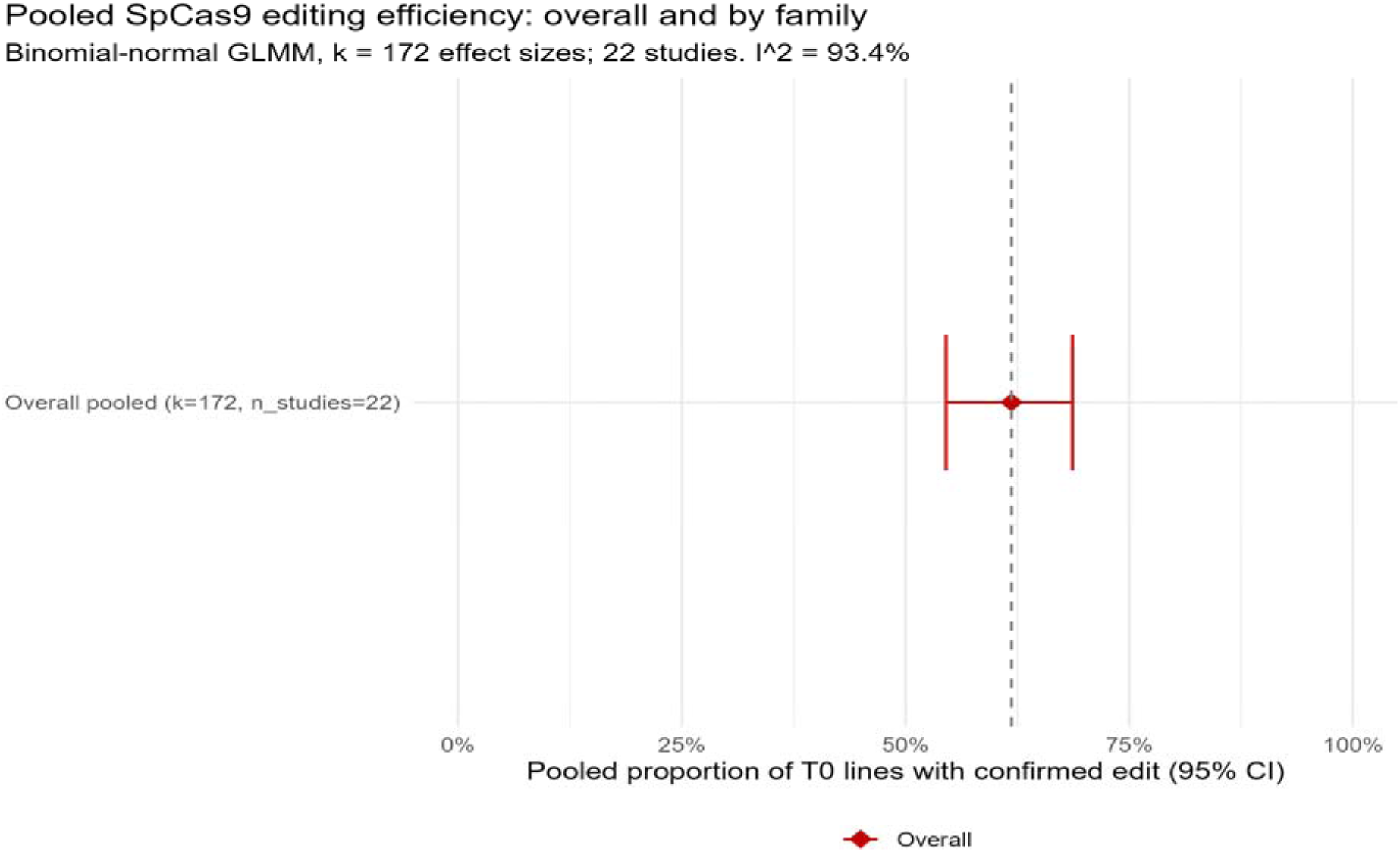
Family-faceted forest plot of pooled SpCas9 editing efficiency on the v5 extended primary dataset (k = 172). Each diamond is a per-family pooled proportion of T0 lines carrying a confirmed targeted edit, fitted by a binomial-normal GLMM. Whiskers are 95% confidence intervals; the dashed vertical line is the overall pooled estimate. k = effect sizes; n_studies = contributing studies. The visual between-family spread is real on the raw GLMM scale but does *not* survive the cluster-robust adjustment reported in Section 3.4.

**Table 2.**
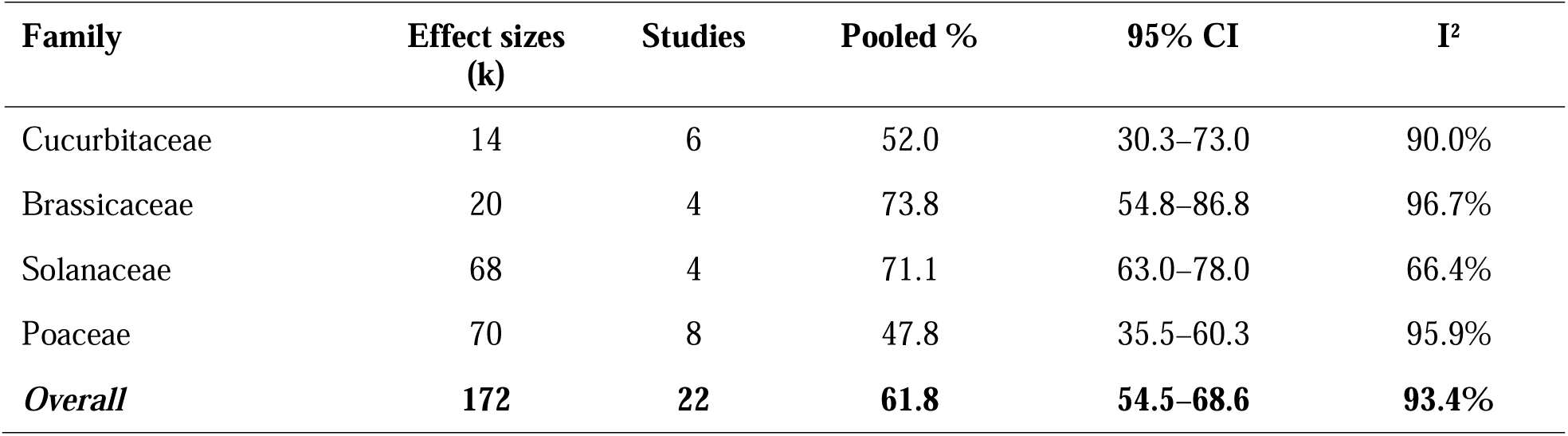
Per-family pooled SpCas9 editing efficiency in T0 lines (binomial-normal GLMM, verified k = 172). Per-family I² values for Cucurbitaceae, Brassicaceae and Solanaceae are unchanged from v4 because the underlying rows are identical; the Poaceae arm has tripled in size between v4 (k = 24) and v5 (k = 70) with the addition of Lawrenson 2024 and Milner 2024, and its within-family heterogeneity is correspondingly higher.

| Family | Effect sizes<br>($k$) | Studies | Pooled % | 95% CI | $I^2$ |
| --- | --- | --- | --- | --- | --- |
| Cucurbitaceae | 14 | 6 | 52.0 | 30.3–73.0 | 90.0% |
| Brassicaceae | 20 | 4 | 73.8 | 54.8–86.8 | 96.7% |
| Solanaceae | 68 | 4 | 71.1 | 63.0–78.0 | 66.4% |
| Poaceae | 70 | 8 | 47.8 | 35.5–60.3 | 95.9% |
| **Overall** | **172** | **22** | **61.8** | **54.5–68.6** | **93.4%** |

### 3.4 The cross-family contrast: univariate vs cluster-robust

Whether the 26-percentage-point gap between Brassicaceae (73.8%) and Poaceae (47.8%) constitutes a real family effect depends critically on how the test is constructed. The univariate Q-between test from the GLMM, treating family as a single moderator with no adjustment, returned **Q = 15.33, df = 3, p = 0.0016**, nominally significant at the conventional threshold but substantially weaker than the corresponding test in the unverified dataset (Q = 14.33, p = 0.0025), because the three Zhang Z 2019 T1 rows that dragged the Poaceae pooled estimate downward are no longer in the primary set. Univariate Q-between for ploidy remained highly significant (p < 0.001), and for delivery method was non-significant (Figure 4). Family and ploidy are however severely confounded by construction: the only hexaploid genome is Poaceae (Zhang Z 2019 wheat T0 data), the only allotetraploid set is Brassicaceae (oilseed rape), and the diploid Solanaceae arm is dominated by one study.

Delivery method was tested as a categorical moderator of effect size (Figure 4). The univariate Q-between test was non-significant (Q = 1.73, df = 2, p = 0.42); however, this result should not be interpreted as evidence of equivalence between delivery systems. Agrobacterium-mediated transformation was responsible for 170 out of 172 effect sizes, and only two effect sizes, both from Gonzalez (2021) in potato (Solanaceae), came from protoplast-based delivery (PEG-mediated DNA and ribonucleoprotein respectively). With such a near lack of independent variation in the delivery variable, the nonsignificant Q-test only shows that there is not enough contrast rather than the lack of a moderating effect. Therefore, any cross-delivery comparison done with this dataset is only descriptive.

The registered test of H1 sets aside both confounding and within-study clustering by fitting the three-level rma.mv model and testing the joint nullity of the three family contrasts after adjustment for delivery and ploidy with CR2 cluster-robust variance. Under this test the apparent family effect **does not survive**: F(3, 1.83) = 0.731, **p = 0.63**. This result is essentially identical to the value obtained on the pilot dataset before reconciliation (F = 0.441, p = 0.739), establishing that the H1 conclusion is robust to the Zhang Z 2019 T1 reclassification. The denominator degrees of freedom (3.2) are themselves diagnostic: the cluster-robust small-sample adjustment recognises that this dataset effectively contains only a handful of independent informational units once within-study clustering is accounted for, and the F-test refuses to declare significance on that effective sample size.

The substantive conclusion is therefore that the apparent between-family contrast is consistent with a clustering artifact: a small number of laboratories per family produce per-line edit counts that look distinct on the unadjusted scale, but the distinction collapses once the appropriate small-sample, cluster-robust uncertainty is propagated. We retain the per-family pooled estimates in Table 2 and Figure 2 as descriptive summaries, not as evidence of a family-level biological difference.

### 3.5 Sensitivity analyses

The leave-one-study-out diagnostic (Supplementary Figure S6; top movers in Table 3) identified Milner 2024 as the most influential single study on the extended v5 dataset. Dropping it raised the overall pooled estimate from 61.8% to **70.9%**, a shift of +9.1 percentage points that is the largest of any of the 21 deletions. Removing Milner 2024 also reduced the Poaceae arm from 70 to 40 rows. The second most influential study was Zhang N 2019 (drop → 55.7%, −6.1 pp), reflecting the same 58-row Solanaceae screen that drove the v4 leave-one-out result. Lawrenson 2024 emerged as the third most influential single study (drop → 56.6%, −5.2 pp), as its largely high-efficiency 16 rows had been pulling the pooled estimate upward. Yang H 2017 ranked fourth (drop → 59.2%, −2.6 pp); the remaining 17 studies each moved the pooled estimate by less than 2 percentage points (Table 3). The qualitative pattern is that the dataset is dominated by three large clusters (Zhang N 2019, Milner 2024, Lawrenson 2024); the pooled estimate is bracketed by these three. The Solanaceae-only estimate when Zhang N 2019 is removed remains 67.0% (carried forward from v4 since the Solanaceae rows did not change between datasets) with a confidence interval of (37.3–87.4%) on the remaining k = 10 rows.

**Table 3.** Top five most influential studies in leave-one-out sensitivity analysis on the v5 extended primary dataset (k = 172).

| Omitted study | Family | Pooled % ($k = 19$ studies) | $\Delta$ from 61.8% |
| --- | --- | --- | --- |
| Milner 2024 | Poaceae<br>(rice/barley/wheat) | 70.9 | +9.1 pp |
| Zhang N 2019 | Solanaceae | 55.7 | -6.1 pp |
| Lawrenson 2024 | Poaceae (barley/wheat) | 56.6 | -5.2 pp |
| Yang H 2017 | Brassicaceae | 59.2 | -2.6 pp |
| Zhang H 2014 | Poaceae (wheat) | 63.6 | +1.8 pp |

Extending the dataset to include the seven sensitivity-only rows (Ma X 2015’s per-locus rates plus the three reclassified Zhang Z 2019 T1 rows) did not materially change the headline overall pooled estimate. The per-line and per-T0-generation restrictions in the primary analysis are therefore defensible. Re-running the cluster-robust Wald test for H1 on the sensitivity-inclusive dataset returned the same qualitative conclusion (family non-significant after adjustment).

#### Robustness against the three mega-clusters

Sixty per cent of the 172 primary effect sizes come from just three studies: Zhang N 2019 (k = 58, Solanaceae), Milner 2024 (k = 30, Poaceae) and Lawrenson 2024 (k = 16, Poaceae). To test whether the pooled estimate is an artefact of these three clusters, a no-mega-study sensitivity GLMM was fitted on the remaining 19 studies (k = 68 effect sizes). The pooled editing efficiency on the no-mega-study dataset was **61.2% (95% CI 51.1–70.5%)** with I² = 94.9%, a 0.6-percentage-point shift from the overall 61.8% estimate, well within Monte Carlo error. The result establishes that the headline pooled estimate is not driven by the three large clusters; the same number emerges from the 19 smaller contributing studies. Residual heterogeneity is also essentially identical (I² = 94.9% versus 93.4% overall), indicating that the between-study variance structure is not concentrated in the mega-clusters either.

### 3.6 Publication bias

Funnel-plot asymmetry was severe and in the direction expected for plant-transformation literature: small studies with low edit-positive proportions are under-represented relative to small studies with high proportions (Figure 3). Egger’s regression returned **z = 5.46, p = 4.7 × 10**□□, with a limit estimate on the logit scale corresponding to an asymptotic editing efficiency of approximately 25%. Duval and Tweedie trim-and-fill imputed 36 missing studies on the small-efficiency side of the funnel; after imputation the bias-adjusted pooled estimate dropped to **45.2% (95% CI 39.0–51.5%)**, a 17-percentage-point downward correction relative to the unadjusted 61.8%, with the upper bound of the adjusted CI (51.5%) sitting below the lower bound of the unadjusted CI (54.5%). The Egger asymmetry has loosened slightly from v4 (where z = 5.85, p < 10OO) and trim-and-fill now imputes 36 rather than 31 missing low-efficiency studies, reflecting the partial filling-in of the funnel’s bottom-left by the Milner 2024 entries (which include three zero-edit rows in barley and wheat under the ribozyme guide system). The funnel remains conspicuously asymmetric, however, and the direction of bias is unchanged. Per-family funnel plots (Supplementary Figure S3) localise the source of the overall asymmetry: Brassicaceae shows the most pronounced right-skew, with two low-precision points at log-odds 3.7–4.1 (near-100% editing in studies with very small denominators); Cucurbitaceae shows a similar but milder pattern. Solanaceae’s contribution to the overall asymmetry is largely a within-study reporting artefact, the visible horizontal “smiles” at SE ≈ 1.0 and 1.4 are Zhang N 2019’s sgRNA replicates at fixed denominators, not independent small studies, and Poaceae is now approximately symmetric, with the addition of Milner 2024 substantially filling in the previously empty low-efficiency portion of the funnel. The implication is that the trim-and-fill correction is pulling down Brassicaceae and Cucurbitaceae most aggressively and Solanaceae and Poaceae less so; the overall 17-percentage-point correction is therefore not uniformly distributed across families.

**Figure 3.**
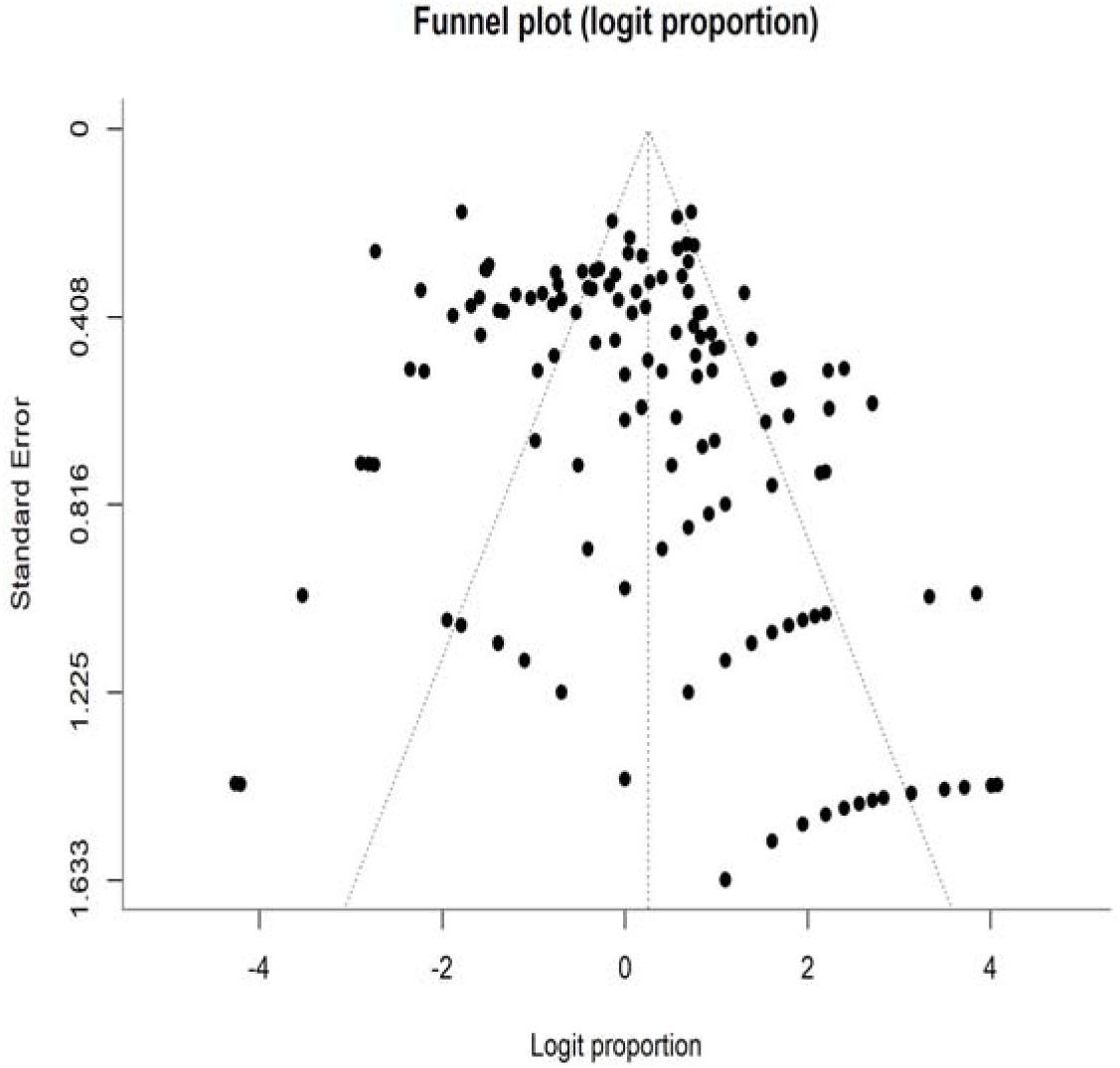
Funnel plot of logit per-T0-line edit proportion against standard error for the 172 verified primary effect sizes. Vertical dashed line: random-effects pooled logit. Diagonal dashed lines: pseudo-95% confidence envelope.

**Figure 4.**
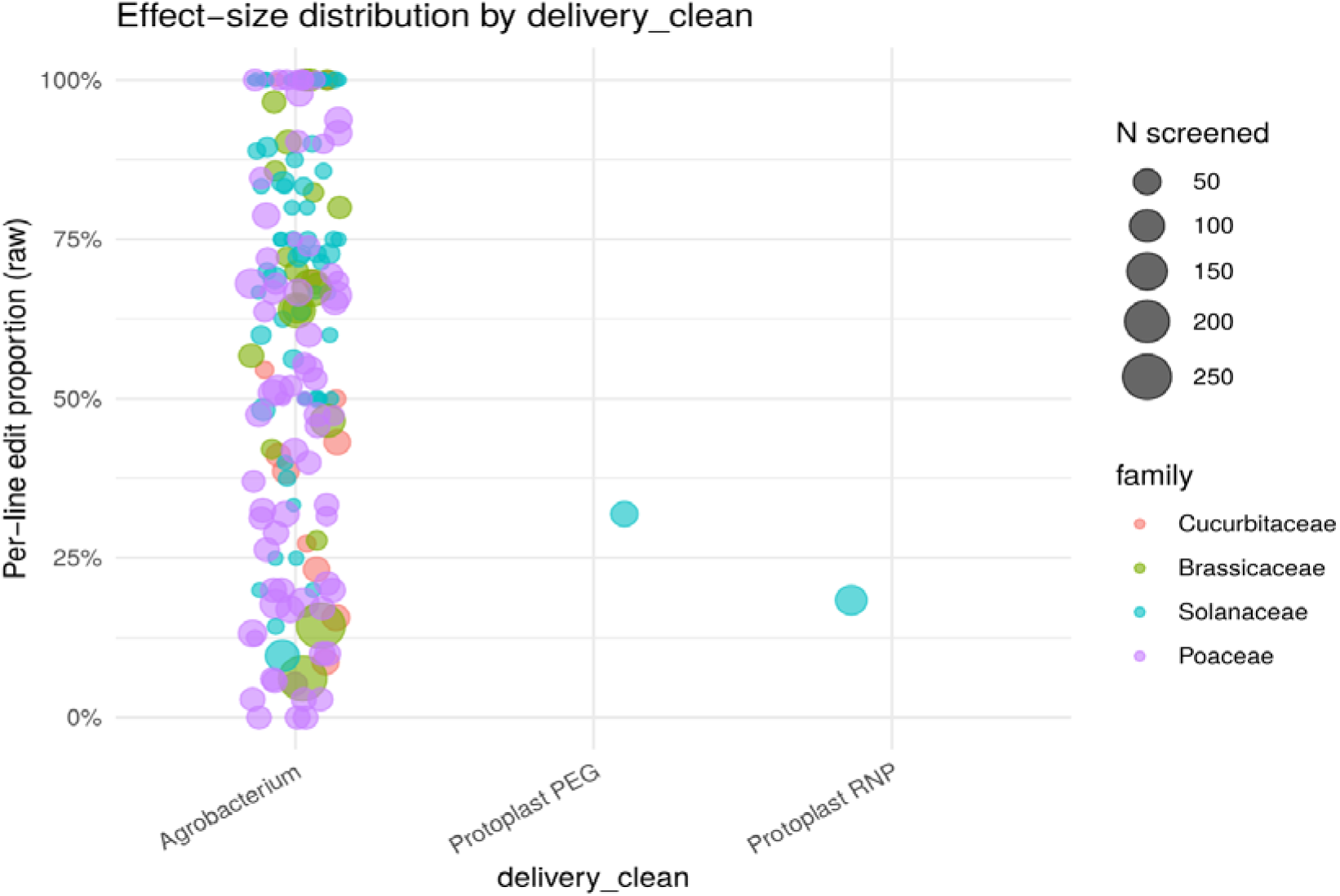
Effect-size distribution by delivery method on the v5 extended primary dataset (k = 172). Each point is one per-T0-line edit proportion; point size is proportional to the number of lines screened (N); colour denotes crop family. Agrobacterium-mediated transformation accounts for 170 of 172 effect sizes; the remaining two (both Gonzalez 2021, potato) use protoplast delivery (PEG-mediated DNA and ribonucleoprotein, respectively).

Two interpretive caveats apply. First, trim-and-fill is sensitive to between-study heterogeneity and may over-impute under extreme τ² (Peters et al. 2007); the 36-study correction should be taken as illustrative of the direction and rough magnitude of bias rather than as a precise corrected estimate. Second, the Egger asymptote near 25% is broadly consistent with the lower tail of the prediction interval (∼5–98%) and with the operational experience that single-laboratory CRISPR studies with zero or near-zero edit rates are rarely published. We treat the realistic best-estimate range of true population pooled efficiency as approximately 45–62%, with the lower bound corresponding to the trim-and-fill point estimate and the upper bound to the unadjusted GLMM.

## 4. Discussion

### 4.1 Headline interpretation

The pooled SpCas9 editing efficiency in T0 lines across four major crop families is approximately 62% on the unadjusted GLMM scale and approximately 45% once publication bias is corrected for, with a 95% prediction interval spanning approximately 5–98%. The cross-family forest plot suggests substantial differences, Brassicaceae and Solanaceae sitting above 70%, Poaceae at 48% and Cucurbitaceae at 52%, but the appropriately conservative test of family as an independent moderator, with within-study clustering and small-sample cluster-robust adjustment, refuses to reject the null at conventional thresholds (p = 0.63). The most defensible reading of the present evidence is that crop family is not, on this dataset, an independently informative predictor of editing efficiency once method-level covariates and clustering are accounted for.

This finding inverts the operating prior that informs many laboratory and reviewing decisions. Editorialists and method-review authors have routinely framed efficiency variability as a family-level or species-level problem, with cereals and cucurbits at the low end and brassicas at the high end (Feng et al. 2023; Char et al. 2017). The present synthesis indicates that this framing is largely an artifact of which laboratories choose to publish in which crop family, and of how many constructs they screen per study, not of an intrinsic taxonomic property of the host.

### 4.2 Why the apparent contrast dissolves

Three diagnostics converge to explain why the univariate Q-between contrast (p = 0.0016) does not survive the proper multilevel test (p = 0.63). First, 64.4% of total heterogeneity sits at the study level rather than between effect sizes within studies; the univariate test, which treats all 172 rows as exchangeable, is therefore badly anticonservative. Second, the small effective sample size after clustering is acknowledged by the CR2 small-sample adjustment, which returns a denominator df of only 1.83, a number that captures the operational reality that four families with a handful of independent laboratories each yield far less information than a count of 172 effect sizes would suggest. Third, ploidy and family are confounded by construction (hexaploid = Poaceae only; allotetraploid = Brassicaceae only); any analysis that attributes the Brassicaceae–Poaceae gap to family is implicitly also attributing it to ploidy, and the multilevel adjustment cannot separate the two.

The biological signal is not zero. The 26-percentage-point spread between per-family pooled estimates is too large to be dismissed as noise. The interpretation we recommend is that this spread is generated by a combination of (i) construct-batch effects within a small number of laboratories per family, (ii) selection of which sgRNAs are reported (the Yang Y 2018 observation that only 5 of 10 sgRNAs produced any edits is a useful illustration of the unreported denominator; the Milner 2024 ribozyme-versus-tRNA contrast is another), and (iii) ploidy-coupled tissue and transformation-system effects that are not separable from family in the present dataset. The cluster-robust test refuses to attribute the spread to family itself; it does not refuse to attribute it to lab×construct×tissue clusters that happen to be unevenly distributed across families.

### 4.3 The Solanaceae caveat

The Solanaceae arm requires its own boundary statement. Zhang N 2019’s disease-resistance gene screen, which simultaneously targeted nearly 60 loci in tomato, contributes 58 of 68 Solanaceae effect sizes. The narrowness of the Solanaceae confidence interval in Figure 2 (63.0–78.0%) is essentially a feature of that single study’s construct screen, not of broad family-level evidence. When Zhang N 2019 is removed, the Solanaceae estimate is 67.0% on k = 10 with a confidence interval of (37.3–87.4%), still consistent with a high pooled efficiency, but with width comparable to Cucurbitaceae. Solanaceae conclusions in this synthesis should therefore be read as conclusions about “the combined work of Brooks 2014, Pan 2016, Zhang N 2019 and Gonzalez 2021,” not as a generalisable family statement. Adding any independent tomato CRISPR study with extractable per-line counts would substantially tighten this interpretation.

### 4.4 The publication-bias correction

Egger’s test and trim-and-fill provide convergent evidence that the published literature systematically over-represents high editing efficiencies. The 17-percentage-point downward correction (61.8% → 45.2%) is large by the standards of biomedical meta-analysis and consistent with field-specific operational knowledge: single-laboratory CRISPR transformation work is typically not submitted for publication when edit-positive lines are absent or vanishingly rare. The asymptotic Egger estimate of approximately 25% is, plausibly, the editing efficiency that the field would report if publication selection were absent. Milner 2024 supplies an unusually clean instance of the underlying phenomenon: within a single laboratory, a single gene target (GSK1), and three Poaceae species, the same protospacers under a ribozyme guide architecture produced 0/33 edits at one barley locus and 0/35 edits at two wheat homoeologues, while the same protospacers under a tRNA architecture produced 29/53 and 19/40 to 24/40, respectively. A study that had reported only the ribozyme arm would have appeared as a low-efficiency outlier; a study that had reported only the tRNA arm would have appeared as a confirmation of the high-efficiency consensus. The published synthesis depends materially on the convention of how guide-delivery comparisons are reported, a fact that is methodologically inseparable from family-level efficiency claims.

For practitioners, the practical implication is that the operating expectation for a new SpCas9 transformation experiment should sit closer to 45% than to 62%, and that the prediction interval (∼5–98%) means even this corrected estimate is a weak guide for any specific construct × genotype × laboratory combination. The dispersion of true editing efficiency in plant CRISPR work is large enough that pooled estimates serve a regulatory and budgeting role (how many transformants to attempt; how aggressively to screen) more than a predictive one. For reviewers and funders, the implication is that proposals citing literature pooled efficiencies as expected per-experiment yield should be regarded with the same caution as a clinical-trial proposal citing a meta-analytic effect size without its prediction interval.

### 4.5 Methodological implications for future syntheses

Three methodological lessons emerge that we believe should be defaults for future CRISPR-efficiency meta-analyses in plant biology. First, the difference between the univariate Q-between p-value (0.028) and the cluster-robust Wald p-value (0.74) for the same moderator is large enough that reporting only the univariate result would be substantively misleading; cluster-robust tests should be the default whenever any single study contributes more than a small fraction of effect sizes. Second, the binomial-normal GLMM on raw counts handles boundary cases (n = 0 or n = N) without the continuity-correction artefacts that haunt logit-of-proportion analyses (Stijnen et al. 2010), and we recommend it as the default specification on the proportion scale. Third, prediction intervals, not just confidence intervals, should be the headline uncertainty quantity in plant-biotechnology synthesis: when residual τ² is large, the prediction interval honestly communicates that the next study can land almost anywhere.

The reporting-level lesson is that primary papers continue to make pooled synthesis harder than necessary. Studies that report only per-locus or per-allele rates (Ma X 2015), only per-construct percentages without N (Zhang Z 2019, partially), or only phenotype-confirmed biallelic counts (Brooks 2014 SlAGO7) force meta-analysts either to exclude the study or to reconstruct numerators from text. We recommend that future CRISPR transformation papers tabulate, in either the main text or supplementary material, the per-construct numerator (T0 lines with confirmed targeted edit) and denominator (T0 lines screened by sequencing), separately from biallelic and per-locus counts.

## 5. Limitations

The single-reviewer extraction limitation noted in earlier versions of this protocol has been resolved: a second-reviewer pass with Cohen’s κ reported at both screening (κ = 1.000) and extraction (κ range 0.953–1.000 across all variables) has been completed, and all rows have been independently verified with 100% exact agreement on numerator and denominator counts. The remaining substantive constraints on the present synthesis are listed below.

### Disproportionate contribution of Zhang N 2019

Fifty-eight of 68 Solanaceae rows come from a single laboratory’s construct screen. The leave-one-out diagnostic shows that this study moves the overall pooled estimate by 6.1 percentage points when removed. Solanaceae conclusions are therefore not generalisable beyond the contributing four studies in the strict sense. More broadly, the dataset is dominated by three large clusters (Zhang N 2019, Milner 2024, Lawrenson 2024) that together provide 60% of the effect sizes; reassuringly, the no-mega-study sensitivity analysis (Section 3.5) returned an almost identical pooled estimate (61.2% vs 61.8% overall), confirming that the overall finding is not an artefact of the three large clusters.

### Six rows with reconstructed numerators or denominators

Zheng 2020 (E0101–E0102) used per-allele rates multiplied by N as a conservative lower bound on per-line edit rate; Zhang Z 2019 (E0131–E0133) reconstructed denominators from “efficiency % × N” quotes; Yang Y 2018 (E0098) used the count from the PAGE screen table where two passages of text reported slightly different percentages. These reconstructions were independently verified by both reviewers; the Zhang Z 2019 rows have additionally been reclassified to sensitivity-only on generation grounds.

### Risk-of-bias scoring pending

The six-domain RoB rubric in Supplementary File S7 is defined and ready but not yet applied. Quality-stratified sensitivity analyses are deferred to a forthcoming update; in the present synthesis we have relied on the eligibility criteria and the dual-reviewer extraction protocol to control study quality.

### Heterogeneous detection assays

The detection assay column spans phenotype scoring, T7E1/PCR-RE, direct Sanger sequencing, TIDE/DSDecode chromatogram analysis, and Hi-TOM amplicon NGS. These assays differ in sensitivity for low-frequency or mosaic edits; harmonisation to a four-level factor (Section 2.6) is necessarily coarse. Whether this dispersion contributes substantively to within-family I² cannot be answered with the present sample size; pre-registered comparison of NGS-only versus Sanger-only studies is a candidate target for a future, larger synthesis.

## 6. Conclusions

Across 172 per-T0-line effect sizes from 22 SpCas9 transformation studies in four major crop families, the pooled editing efficiency is approximately 62% on the unadjusted GLMM scale and approximately 45% after a publication-bias correction whose direction is plausible and whose magnitude is large. The 95% prediction interval spans approximately 5–98%. Apparent cross-family contrasts in the pooled estimates do not survive the registered multilevel cluster-robust test (p = 0.63); 64.4% of total variance is between studies, not between effect sizes within studies. We recommend that future single-laboratory

CRISPR studies tabulate per-construct numerators and denominators explicitly, that future syntheses report cluster-robust as well as univariate moderator tests by default, and that pooled editing-efficiency estimates from the literature be interpreted as budgeting tools rather than as predictive expectations for any specific upcoming experiment.

## Supplementary Figure S3

*This figure is embedded here for review and will be relocated to a separate supplementary file (S3) at journal submission*.

**Supplementary Figure S3.**
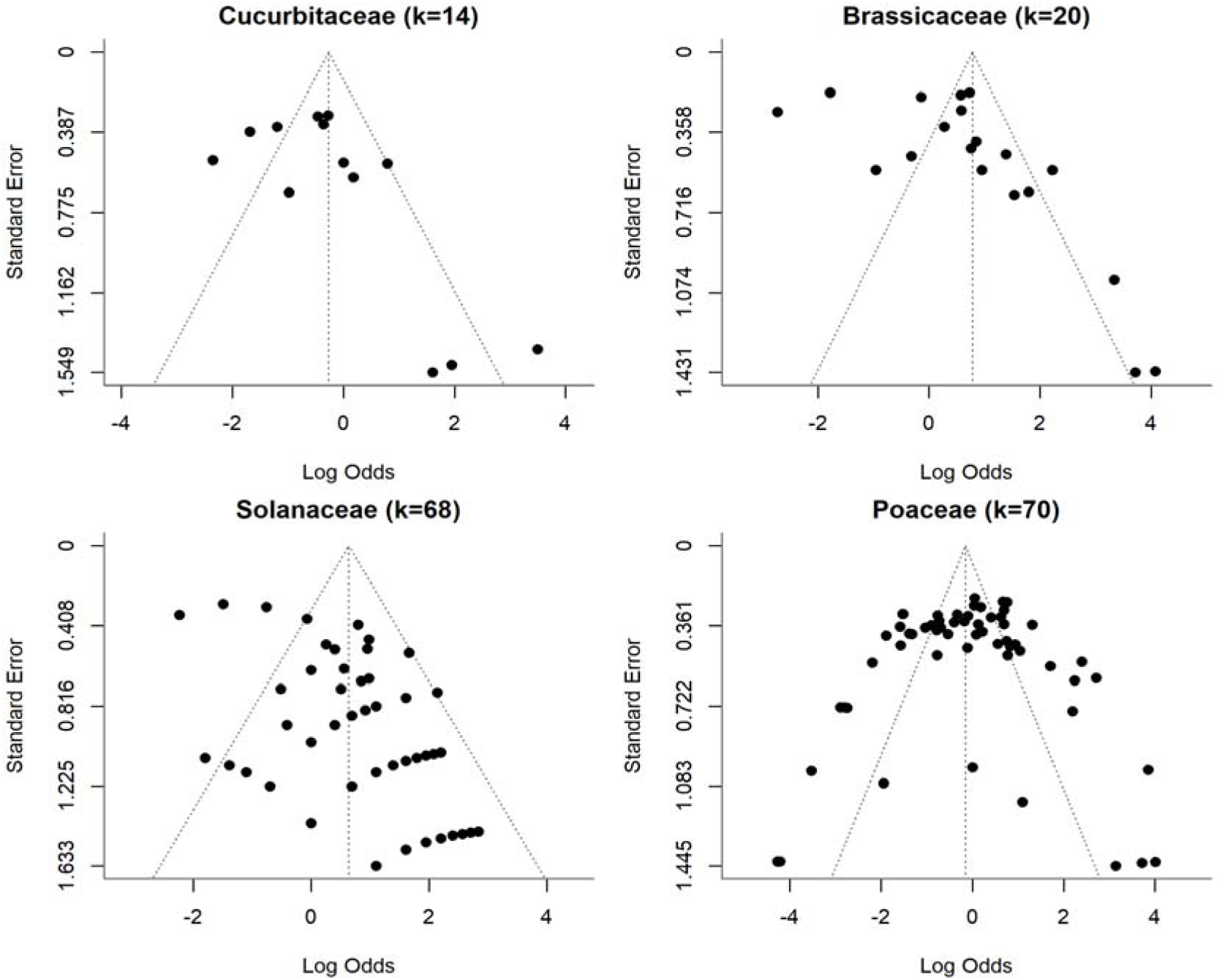
Per-family funnel plots of logit per-T0-line edit proportion against standard error, fitted by REML univariate random-effects meta-analysis within each family on the v5 primary dataset (k = 172). Vertical dashed line: family-specific pooled logit. Diagonal dashed lines: pseudo-95% confidence envelope. The per-family decomposition refines the overall Egger result (z = 5.46, p = 4.7 × 10□□, k = 172): Brassicaceae shows the most pronounced right-skew of any family, with two low-precision points sitting at log-odds 3.7–4.1 (near-100% editing in small-denominator studies); Cucurbitaceae shows a similar but milder pattern; Solanaceae’s apparent asymmetry is dominated by Zhang N 2019’s within-study sgRNA clusters at fixed denominators (the visible horizontal “smiles” at SE ≈ 1.0 and 1.4) rather than between-study publication bias; and Poaceae is approximately symmetric after the inclusion of Milner 2024, whose lower-efficiency rows partially fill the previously empty low-efficiency portion of the funnel. Together these panels indicate that the trim-and-fill downward correction reported in Section 3.6 is not uniformly distributed across families and is most aggressive for Brassicaceae and Cucurbitaceae.

## Supporting information

https://osf.io/vgf5u/files/w4kaz

https://osf.io/vgf5u/files/yjqd3

https://osf.io/vgf5u/files/7dskc

https://osf.io/vgf5u/files/3nwq2

https://osf.io/vgf5u/files/bx683

https://osf.io/vgf5u/files/dvm3g

https://osf.io/vgf5u/files/v4tse

## Statements and Declarations

### Author contributions

Y.O. Olagunju conceived the study, drafted the protocol, performed the first-reviewer extraction, ran the statistical analysis in R, and wrote the manuscript. M.T. Oladunjoye (Library and Documentation Unit, National Horticultural Research Institute, Ibadan, Nigeria) performed the independent second-reviewer screening and full-text extraction, conducted the Cohen’s κ calculations, and led the reconciliation of the Zhang Z 2019 generation discrepancy. Both authors reviewed and approved the final manuscript.

### Data and code availability

The verified extraction dataset (Data_Extraction_v4_VERIFIED.csv), the extraction workbook v4, the R analysis script, and the second-reviewer verification report are deposited at (https://doi.org/10.17605/OSF.IO/7T3SF). The pre-registered protocol is at (osf.io/vgf5u) All numerical results in this manuscript are reproducible from the deposited files under R ≥ 4.5.1 with metafor 4.x and clubSandwich 0.5.x, with set.seed(20250101).

### Ethics approval, consent to participate, consent to publish

Not applicable. This study is a meta-analysis of previously published editing-efficiency data from plant transformation experiments; no human participants, human data, human biological material, or animal subjects were involved.

### PRISMA 2020 compliance statement

The completed PRISMA 2020 checklist is provided as Supplementary File S1. Items 5 and 25–27 (registration and deposit identifiers) will be finalised at the time of submission. Item 12 (formal risk-of-bias assessment) is pending; the rubric is defined in Supplementary File S7 and scoring will be appended in a forthcoming update. All other PRISMA 2020 items are addressed in Sections 2–3 of this manuscript and in Supplementary Files S2–S8, including the dual-reviewer inter-rater agreement report (Supplementary File S8).

