## Supplementary material for "Editing Efficiency Across Crop Families: A Systematic Review and Meta-Analysis of CRISPR/SpCas9 Knockout Outcomes in Cucurbitaceae, Brassicaceae, Solanaceae and Poaceae": https://osf.io/vgf5u/files/w4kaz

**Supplementary File S1**

**PRISMA 2020 Reporting Checklist**

*CRISPR/SpCas9 editing efficiency across four crop families: a systematic review and meta-analysis (Cucurbitaceae, Brassicaceae, Solanaceae, Poaceae)*

This checklist follows the PRISMA 2020 reporting standard (Page et al. 2021, *BMJ* 372:n71). Each row indicates the section, item number, the checklist requirement (paraphrased from the source), and the location in the manuscript and supplementary materials where the item is reported. Where a checklist item is not yet completed, this is stated explicitly.

### **Item-by-item compliance**

| **Section** | **Item #** | **Checklist requirement (paraphrased)** | **Location reported** |
| --- | --- | --- | --- |
| **TITLE** | **1** | Identify the report as a systematic review. | Title page; running head |
| **ABSTRACT** | **2** | See PRISMA-2020 Abstract checklist (PRISMA 2020 statement, Box). | Abstract (p.1–2) |
| **INTRODUCTION** | **3** | Describe the rationale for the review in the context of existing knowledge. | 1 Introduction, p.3–4 (paragraphs 1–2) |
|  | **4** | Provide an explicit statement of the objective(s) or question(s) the review addresses. | 1 Introduction, p.4 (final paragraph): O1–O3 + H1 |
| **METHODS** | **5** | Specify the inclusion and exclusion criteria for the review and how studies were grouped for the syntheses. | 2.2 Eligibility criteria, p.5 |
|  | **6** | Specify all databases, registers, websites, organisations, reference lists and other sources searched or consulted to identify studies; specify the date when each source was last searched or consulted. | 2.3 Information sources and search, p.5; Supplementary File S2 (Boolean strings + dates) |
|  | **7** | Present the full search strategies for all databases, registers and websites, including any filters and limits used. | Supplementary File S2 (this package) |
|  | **8** | Specify the methods used to decide whether a study met the inclusion criteria of the review, including how many reviewers screened each record and each retrieved report, whether they worked independently, and, if applicable, details of automation tools used. | 2.4 Study selection and data extraction (dual-reviewer), p.5–6; Supplementary File S8 (verification report) |
|  | **9** | Specify the methods used to collect data from reports, including how many reviewers collected data from each report, whether they worked independently, any processes for obtaining or confirming data from study investigators, and, if applicable, details of automation tools used. | 2.4, p.5–6; Supplementary File S8 |
|  | **10a** | List and define all outcomes for which data were sought. Specify whether all results that were compatible with each outcome domain in each study were sought (e.g. for all measures, time points, analyses), and if not, the methods used to decide which results to collect. | 2.2 (primary outcome: per-T0-line edit proportion); 2.4 (primary_outcome_flag column described in Supplementary File S4) |
|  | **10b** | List and define all other variables for which data were sought (e.g. participant and intervention characteristics, funding sources). Describe any assumptions made about any missing or unclear information. | Supplementary File S4 (moderator harmonisation rules); Codebook (Supplementary File S5) |
|  | **11** | Specify the methods used to assess risk of bias in the included studies, including details of the tool(s) used, how many reviewers assessed each study and whether they worked independently, and, if applicable, details of automation tools used. | 2.5 Risk of bias assessment, p.6; Supplementary File S7 (risk-of-bias scoring matrix — pending) |
|  | **12** | Specify for each outcome the effect measure(s) (e.g. risk ratio, mean difference) used in the synthesis or presentation of results. | 2.6 Effect measure and synthesis methods, p.6 (logit proportion of T0 lines with confirmed edit) |
|  | **13a** | Describe the processes used to decide which studies were eligible for each synthesis (e.g. tabulating the study intervention characteristics and comparing against the planned groups for each synthesis). | 2.4, p.5–6; Figure 1 PRISMA flow diagram |
|  | **13b** | Describe any methods required to prepare the data for presentation or synthesis, such as handling of missing summary statistics, or data conversions. | 2.4, p.6 (handling of zero numerators; reconstruction of integer counts where only percentages reported) |
|  | **13c** | Describe any methods used to tabulate or visually display results of individual studies and syntheses. | 2.7, p.6–7; Figures 2 (forest), 3 (funnel), 4 (delivery bubble); Supplementary Figure S3 (per-family funnels), Supplementary Figure S6 (leave-one-out) |
|  | **13d** | Describe any methods used to synthesise results and provide a rationale for the choice(s). If meta-analysis was performed, describe the model(s), method(s) to identify the presence and extent of statistical heterogeneity, and software package(s) used. | 2.6, p.6 (binomial-normal GLMM via rma.glmm; rma.mv with clubSandwich CR2 robust variance); R 4.5.1, metafor 4.x, clubSandwich 0.5.x |
|  | **13e** | Describe any methods used to explore possible causes of heterogeneity among study results (e.g. subgroup analysis, meta-regression). | 2.7 Subgroup and moderator analyses, p.7 (family, delivery_clean, ploidy_clean, sgRNA_promoter_clean, target_class_clean, detection_assay_clean) |
|  | **13f** | Describe any sensitivity analyses conducted to assess robustness of the synthesised results. | 2.7, p.7 (leave-one-out; no-mega-study; sensitivity-inclusive dataset; per-locus inclusion sensitivity) |
|  | **14** | Describe any methods used to assess risk of bias due to missing results in a synthesis (arising from reporting biases). | 2.8 Publication bias, p.7 (Egger’s regression; Duval and Tweedie trim-and-fill) |
|  | **15** | Describe any methods used to assess certainty (or confidence) in the body of evidence for an outcome. | 2.5, p.6; not formally GRADE-assessed (rationale in 5 Limitations) |
| **RESULTS** | **16a** | Describe the results of the search and selection process, from the number of records identified in the search to the number of studies included in the review, ideally using a flow diagram. | 3.1, p.7–8; Figure 1 PRISMA flow diagram |
|  | **16b** | Cite studies that might appear to meet the inclusion criteria, but which were excluded, and explain why they were excluded. | Supplementary File S5 (excluded studies log; reasons summarised in Figure 1) |
|  | **17** | Cite each included study and present its characteristics. | Table 1, p.8; Supplementary File S5 (per-study metadata) |
|  | **18** | Present assessments of risk of bias for each included study. | Supplementary File S7 (pending; rubric described in 2.5) |
|  | **19** | For all outcomes, present, for each study: (a) summary statistics for each group (where appropriate) and (b) an effect estimate and its precision (e.g. confidence/credible interval), ideally using structured tables or plots. | Figure 2 forest plot; underlying data in Supplementary File S5 |
|  | **20a** | For each synthesis, briefly summarise the characteristics and risk of bias among contributing studies. | 3.1–3.2, p.7–9 |
|  | **20b** | Present results of all statistical syntheses conducted. If meta-analysis was done, present for each the summary estimate and its precision (e.g. confidence/credible interval) and measures of statistical heterogeneity. If comparing groups, describe the direction of the effect. | 3.2–3.3, Tables 2–3, Figures 2–4 |
|  | **20c** | Present results of all investigations of possible causes of heterogeneity among study results. | 3.4 H1 multilevel cluster-robust test, p.10; 3.5 Sensitivity analyses, p.11–12 |
|  | **20d** | Present results of all sensitivity analyses conducted to assess the robustness of the synthesised results. | 3.5, p.11–12 (leave-one-out, no-mega-study robustness, sensitivity-inclusive dataset) |
|  | **21** | Present assessments of risk of bias due to missing results (arising from reporting biases) for each synthesis assessed. | 3.6 Publication bias, p.13–14; Figure 3 funnel plot; Supplementary Figure S3 (per-family funnel decomposition) |
|  | **22** | Present assessments of certainty (or confidence) in the body of evidence for each outcome assessed. | 3.6–3.7; cumulative confidence narrative in 4 |
| **DISCUSSION** | **23a** | Provide a general interpretation of the results in the context of other evidence. | 4.1–4.2, p.15–16 |
|  | **23b** | Discuss any limitations of the evidence included in the review. | 4.3 + 5 Limitations, p.17–18 |
|  | **23c** | Discuss any limitations of the review processes used. | 5 Limitations, p.17–18 (search dates; AI-assisted extraction of Lawrenson/Milner pending second-reviewer verification; risk-of-bias scoring pending) |
|  | **23d** | Discuss implications of the results for practice, policy, and future research. | 4.4 Implications for practice + 6 Conclusions |
| **OTHER INFORMATION** | **24a** | Provide registration information for the review, including register name and registration number, or state that the review was not registered. | 2.1, p.5; OSF registration ID (placeholder to be populated post-submission) |
|  | **24b** | Indicate where the review protocol can be accessed, or state that a protocol was not prepared. | 2.1, p.5; protocol deposited at OSF project ID (placeholder) |
|  | **24c** | Describe and explain any amendments to information provided at registration or in the protocol. | 2.1; v5 extension (Lawrenson + Milner addendum) documented in Supplementary File S8; reclassification of three Zhang Z 2019 T1 rows from primary to sensitivity is documented in 2.4 |
|  | **25** | Describe sources of financial or non-financial support for the review, and the role of the funders or sponsors in the review. | Funding statement (title page) |
|  | **26** | Declare any competing interests of review authors. | Conflicts of interest statement (title page) |
|  | **27** | Report which of the following are publicly available and where they can be found: template data collection forms; data extracted from included studies; data used for all analyses; analytic code; any other materials used in the review. | 2.1, p.5; full deposit at OSF project; analytical R pipeline and dataset are accessible via the OSF DOI |
