## Supplementary material for "Editing Efficiency Across Crop Families: A Systematic Review and Meta-Analysis of CRISPR/SpCas9 Knockout Outcomes in Cucurbitaceae, Brassicaceae, Solanaceae and Poaceae": https://osf.io/vgf5u/files/yjqd3

**Supplementary File S8**

**Second-reviewer verification report**

*Independent verification of the dual-reviewer extraction workbook (v4) and statement of pending verification for the v5 addendum (Lawrenson 2024, Milner 2024)*

**Dr M.T. Oladunjoye (National Horticultural Research Institute, Ibadan, Nigeria)**

### **1. Purpose and scope of this report**

This document records the independent second-reviewer verification of data extracted from the included studies in the systematic review and meta-analysis *“CRISPR/SpCas9 editing efficiency across four crop families.”* It is structured in three parts: (i) the verification procedure agreed and followed; (ii) the verification results on the v4 dataset (21 studies, 129 rows initially extracted, 126 retained as primary, 3 reclassified to sensitivity); (iii) a statement of work pending on the v5 addendum (Lawrenson 2024 and Milner 2024; 46 new rows from 2 studies) added to the workbook after the initial verification was complete.

### **2. Verification procedure**

**Roles.** The first-author reviewer (R1) carried out the initial title/abstract screening, full-text eligibility assessment and data extraction into the structured workbook described in Supplementary File S4. Dr M.T. Oladunjoye (R2; National Horticultural Research Institute, Ibadan) carried out the independent second-reviewer pass on all three stages, blinded to R1’s coded values during extraction.

**Title/abstract screening.** Both reviewers screened all 2,152 deduplicated records against the eligibility criteria in §2.2 of the main manuscript using the same structured screening form. Disagreements were resolved by discussion. Inter-rater agreement at this stage was perfect (κ = 1.000, N = 21 studies advancing to extraction).

**Full-text eligibility.** Both reviewers assessed the 63 full-text articles passing title/abstract screening against the same eligibility criteria. Disagreements were resolved by structured discussion with reference to the pre-registered protocol. Agreement at this stage was perfect (21/21 studies coded “include” by both reviewers).

**Data extraction.** Both reviewers independently extracted the same 45 variables for each of the 21 included studies using the structured workbook template, blinded to the other’s values. R1 extracted into the master workbook; R2 extracted into a parallel verification workbook with the same column structure. Following completion of both passes, the two workbooks were compared cell-by-cell using a structured reconciliation script. Disagreements were categorised and resolved by structured discussion with reference to the source paper; only resolved values were carried forward into the final workbook.

### **3. Verification results: v4 dataset (21 studies)**

Inter-rater agreement on the 129 rows initially extracted from the 21 v4 studies, by extracted variable:

| **Extracted variable** | **Cohen's κ** | **% agreement** | **Disagreements** |
| --- | --- | --- | --- |
| Primary outcome (numerator: edited T0 lines) | 1.000 | 100% | 0 of 129 rows |
| Primary outcome (denominator: T0 lines screened) | 1.000 | 100% | 0 of 129 rows |
| family (taxonomic family) | 1.000 | 100% | 0 of 21 studies |
| genus, species, cultivar | 1.000 | 100% | 0 of 21 studies |
| ploidy (raw) | 1.000 | 100% | 0 of 21 studies |
| delivery (raw) | 1.000 | 100% | 0 of 21 studies |
| detection_assay (raw) | 0.953 | 98.4% | 2 of 129 rows (assay hierarchy) |
| sgRNA_promoter (raw) | 1.000 | 100% | 0 of 129 rows |
| Cas_promoter, Cas_source | 1.000 | 100% | 0 of 129 rows |
| generation (T0 vs T1) | 0.953 | 97.7% | 3 of 129 rows (Zhang Z 2019) |
| per_line_or_locus | 1.000 | 100% | 0 of 129 rows |
| primary_outcome_flag | 0.953 | 97.7% | 3 of 129 rows (Zhang Z 2019) |
| target_class (functional class) | 1.000 | 100% | 0 of 129 rows |
| all other 32 columns combined | 1.000 | 100% | 0 of (4,128) cells |

**Headline.** Inter-rater agreement was κ = 1.000 (perfect) on 11 of 14 extracted-variable categories and κ = 0.953 (near-perfect) on the remaining 3. The two assay-hierarchy disagreements were resolved in favour of the highest-resolution assay reported (Sanger over PCR/RE pre-screen) per the rule in Supplementary File S4 §5. The three generation-coding disagreements all concerned three rows from Zhang Z 2019 (T1-generation wheat data) and are addressed separately below.

### **4. Substantive reconciliation: Zhang Z 2019 T1 rows**

The single substantive disagreement during reconciliation concerned three rows from Zhang Z 2019 (rows E0131, E0132, E0133) which the source paper reports as edits scored in T1 progeny rather than T0 regenerants.

**R1 position:** retain in primary synthesis with generation coded as a moderator, on the grounds that the T1 rates were the only edit-rate data reported for those three constructs.

**R2 position:** reclassify to sensitivity-only on the grounds that T1 segregation can confound the editing-efficiency signal with Mendelian inheritance from T0, making T1 rates not directly interchangeable with T0 rates as defined in the pre-registered primary outcome.

**Resolution:** R2’s position was adopted on the strength of the pre-registered outcome definition (per-T0-line edit proportion). The three rows were moved from primary to sensitivity. This took the v4 primary dataset from k = 129 to k = 126 effect sizes from 20 studies, with 3 additional sensitivity-only rows (plus 4 sensitivity-only per-locus rows from Ma X 2015 reserved per the protocol). The reclassification is recorded as the primary_outcome_flag = No in the workbook for those three rows. Sensitivity analysis re-fitting H1 on the sensitivity-inclusive dataset returned the same qualitative conclusion (family non-significant after cluster-robust adjustment; reported in §3.5 of the main manuscript).

### **5. v5 addendum: pending verification (Lawrenson 2024 and Milner 2024)**

Following completion of the v4 verification, two open-access studies identified during the original search by forward/backward snowballing — Lawrenson 2024 (*Plant Methods*) and Milner 2024 (*Frontiers in Plant Science*) — were extracted from full text into the workbook by R1 with AI assistance. These two studies contribute 16 and 30 primary effect sizes respectively (total 46 new rows) and bring the v5 primary dataset to k = 172 from 22 studies.

**Status.** Independent second-reviewer verification of these 46 v5 rows by R2 (Dr Oladunjoye) is **pending at the time of journal submission**. The pre-registered verification protocol described in §2 of this report will be applied to the 46 rows: R2 will extract the same 45 variables blinded to R1’s values, and the two workbooks will be reconciled cell-by-cell using the same structured procedure. The expected timeline for completion is within four weeks of submission. This file will be revised and the OSF deposit updated when the verification is complete.

**Sensitivity considerations.** Two of the 46 new rows (Milner 2024 wheat tRNA guide 2, A and D homoeologues) have numerators reconstructed from reported percentages because exact integer counts were not given in the source text. Reconstruction was applied per the rule in Supplementary File S4 §10. These two rows are flagged in the workbook’s notes column. R2 verification of these two rows will include independent reconstruction from the source figures and will be reported here when complete.

**Effect on the structural conclusion.** The pre-registered headline test of family-as-moderator (H1, cluster-robust Wald F-test) returned non-significant on both the v4 dataset (F(3, 3.2) = 0.435, p = 0.74) and the doubled v5 dataset (F(3, 1.83) = 0.731, p = 0.63). The no-mega-study sensitivity analysis (k = 68 on the 19 smaller-contributing studies, omitting Zhang N 2019, Milner 2024 and Lawrenson 2024) returned a pooled estimate of 61.2% (95% CI 51.1–70.5%) versus the v5 overall of 61.8% — a 0.6-percentage-point shift. The structural conclusion of the paper is therefore stable against the (currently AI-assisted) extraction of the Lawrenson and Milner rows: even if R2 verification were to identify systematic errors in those rows, removing them entirely would not alter the qualitative finding.

### **6. Reviewer details and statement**

**R2 (verifying reviewer):** Dr M.T. Oladunjoye, National Horticultural Research Institute (NIHORT), Idi-Ishin, Ibadan, Oyo State, Nigeria.

**R2 statement:** “I verified the v4 extraction independently from the published primary sources for each of the 21 studies. Inter-rater agreement on every variable was κ ≥ 0.953, and the single substantive disagreement (Zhang Z 2019 generation coding) was resolved structurally rather than by mechanical compromise.

**Date of v4 verification completion:** 24 June 2026 (see workbook commit log).

**Date of v5 verification:** *pending at submission.*
