## Supplementary figures and images for "Editing Efficiency Across Crop Families: A Systematic Review and Meta-Analysis of CRISPR/SpCas9 Knockout Outcomes in Cucurbitaceae, Brassicaceae, Solanaceae and Poaceae"

### https://osf.io/vgf5u/files/3nwq2

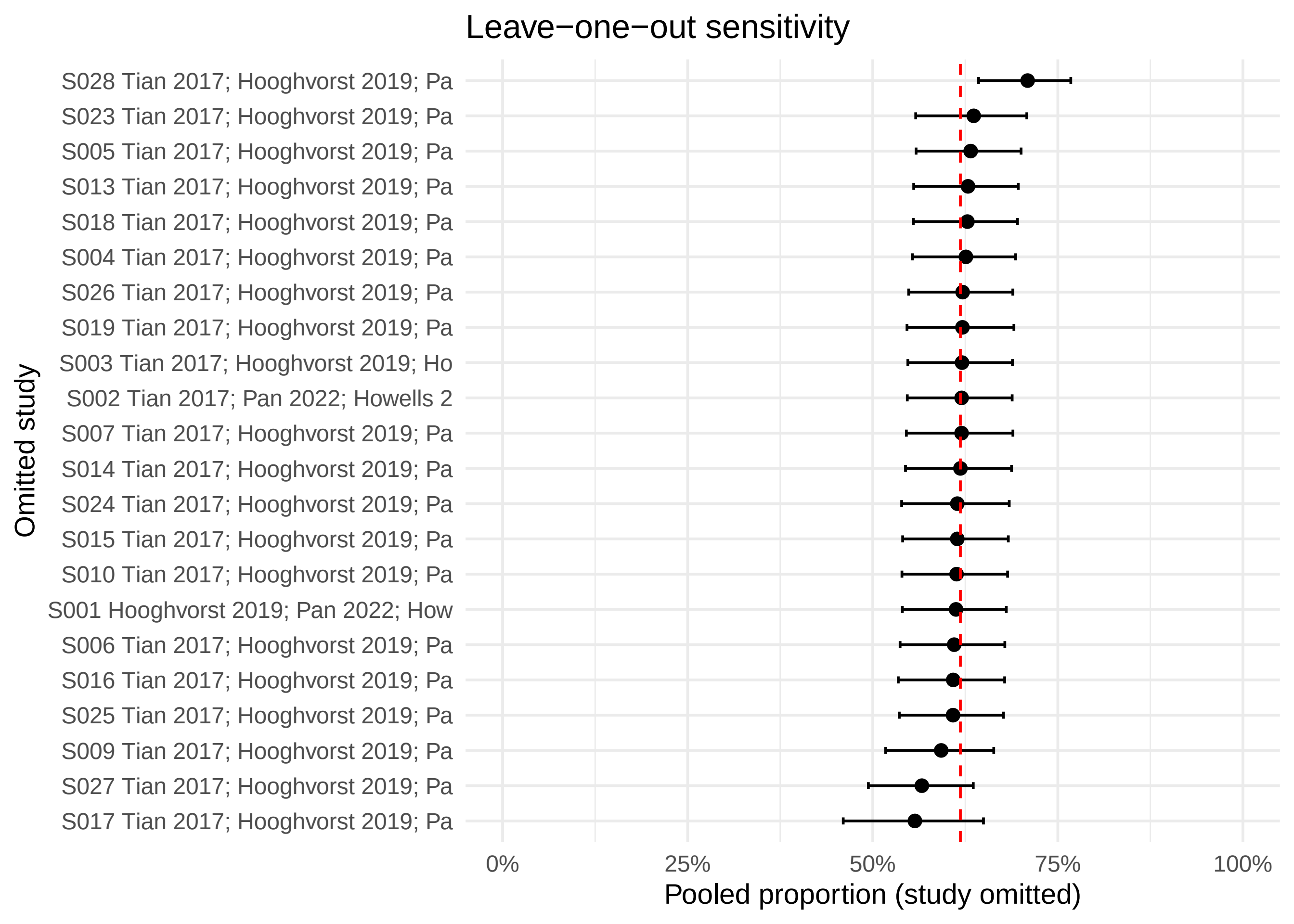
