## Supplementary material for "Editing Efficiency Across Crop Families: A Systematic Review and Meta-Analysis of CRISPR/SpCas9 Knockout Outcomes in Cucurbitaceae, Brassicaceae, Solanaceae and Poaceae": https://osf.io/vgf5u/files/bx683

**Supplementary File S2**

**Database-specific Boolean search strings and per-database hit counts**

*Companion to: "Editing efficiency across crop families: a systematic review and meta-analysis of CRISPR/SpCas9 knockout outcomes in Cucurbitaceae, Brassicaceae, Solanaceae and Poaceae" — §2.3 Information sources and search*

### **1. Overview**

This document reports the verbatim Boolean search strings executed against each of the seven bibliographic databases identified in §2.3 of the main manuscript, together with the per-database record counts at each stage of the PRISMA 2020 flow. Final search date: **23 June 2026**. All searches were limited to peer-reviewed primary-research articles published in English; no date restriction was applied at the database level (date filtering was handled at screening). Forward and backward snowballing was performed on every included study, and the resulting additional records are reported separately in §11.

The four conceptual blocks underlying every database string are:

**•** Concept 1 — CRISPR/SpCas9 nuclease: CRISPR, Cas9, SpCas9, “Streptococcus pyogenes Cas9”, and MeSH/CABI subject headings where available.

**•** Concept 2 — Genome editing / targeted mutagenesis: “genome editing”, “gene editing”, “targeted mutagenesis”, knockout, “loss-of-function”, “editing efficiency”, “mutation frequency”.

**•** Concept 3 — The four target plant families (and key genera/species terms): Cucurbitaceae (cucumber, melon, watermelon, squash, pumpkin, gourd, Cucumis, Citrullus, Cucurbita, Lagenaria); Brassicaceae (Brassica, Arabidopsis, oilseed, rapeseed, canola, mustard, cabbage, broccoli, cauliflower); Solanaceae (tomato, potato, tobacco, pepper, eggplant, Solanum, Nicotiana, Capsicum); Poaceae (rice, wheat, maize, barley, sorghum, millet, Oryza, Triticum, Zea, Hordeum).

**•** Concept 4 — T0/regenerated-plant outcome layer: “T0 lines”, “T0 plants”, “T0 generation”, “regenerated”, “transformed lines”. Applied as a soft filter where database syntax allows; otherwise relaxed to maximise recall at the database stage and applied at title/abstract screening.

Boolean syntax differs across databases; each section below provides the native string for that platform together with the field-tag conventions used.

### **2. PubMed / MEDLINE**

**Platform:** PubMed (NCBI) — https://pubmed.ncbi.nlm.nih.gov/. **Field tags used:** [MeSH] (Medical Subject Heading), [TIAB] (Title/Abstract), [TW] (Text Word). **Truncation:** * (zero or more characters). **Phrase search:** double quotation marks.

##### ***Boolean string (run as a single query)***

```
(
  "CRISPR-Cas Systems"[MeSH] OR "CRISPR-Cas9"[TIAB] OR
  CRISPR[TIAB] OR Cas9[TIAB] OR SpCas9[TIAB] OR
  "Streptococcus pyogenes Cas9"[TIAB]
)
AND
(
  "Gene Editing"[MeSH] OR "genome editing"[TIAB] OR "gene editing"[TIAB] OR
  "targeted mutagenesis"[TIAB] OR "site-directed mutagenesis"[TIAB] OR
  knockout*[TIAB] OR "knock-out"[TIAB] OR "loss-of-function"[TIAB] OR
  "editing efficiency"[TIAB] OR "mutation frequency"[TIAB] OR
  "mutagenesis efficiency"[TIAB]
)
AND
(
  Cucurbitaceae[MeSH] OR Brassicaceae[MeSH] OR Solanaceae[MeSH] OR Poaceae[MeSH] OR
  cucumber[TIAB] OR melon[TIAB] OR watermelon[TIAB] OR squash[TIAB] OR
  pumpkin[TIAB] OR cucurbit*[TIAB] OR gourd[TIAB] OR
  Cucumis[TIAB] OR Citrullus[TIAB] OR Cucurbita[TIAB] OR Lagenaria[TIAB] OR
  Brassica[TIAB] OR Arabidopsis[TIAB] OR oilseed[TIAB] OR rapeseed[TIAB] OR
  canola[TIAB] OR mustard[TIAB] OR cabbage[TIAB] OR broccoli[TIAB] OR
  cauliflower[TIAB] OR Raphanus[TIAB] OR
  tomato[TIAB] OR potato[TIAB] OR tobacco[TIAB] OR pepper[TIAB] OR
  eggplant[TIAB] OR Solanum[TIAB] OR Nicotiana[TIAB] OR Capsicum[TIAB] OR
  rice[TIAB] OR wheat[TIAB] OR maize[TIAB] OR corn[TIAB] OR barley[TIAB] OR
  sorghum[TIAB] OR millet[TIAB] OR cereal*[TIAB] OR
  Oryza[TIAB] OR Triticum[TIAB] OR "Zea mays"[TIAB] OR Hordeum[TIAB]
)
AND
(
  plant*[TIAB] OR transformation[TIAB] OR transformant*[TIAB] OR
  regenerat*[TIAB] OR T0[TIAB] OR "T0 lines"[TIAB] OR "T0 plants"[TIAB]
)
AND English[Language]
```

**Filters:** Language = English. No date filter at database level (date range handled at screening). No publication-type exclusions; “not SpCas9”, “protoplast-only”, and “review” exclusions were applied at full-text assessment, not at the database stage.

### **3. Web of Science Core Collection**

**Platform:** Web of Science (Clarivate) — webofscience.com. **Indexes searched:** SCI-EXPANDED, SSCI, A&HCI, CPCI-S, CPCI-SSH, BKCI-S, BKCI-SSH, ESCI. **Field tags used:** TS= (Topic; title, abstract, author keywords, Keywords Plus). **Truncation:** * (zero or more) and $ (zero or one). **Phrase search:** double quotation marks.

##### ***Boolean string***

```
TS=(
  (CRISPR OR Cas9 OR SpCas9 OR "Streptococcus pyogenes Cas9")
  AND
  ("genome editing" OR "gene editing" OR "targeted mutagenesis" OR
   "site-directed mutagenesis" OR knockout* OR "loss-of-function" OR
   "editing efficiency" OR "mutation frequency" OR "mutagenesis efficiency")
  AND
  (Cucurbitaceae OR Brassicaceae OR Solanaceae OR Poaceae OR
   cucumber OR melon OR watermelon OR squash OR pumpkin OR cucurbit* OR
   Cucumis OR Citrullus OR Cucurbita OR Lagenaria OR
   Brassica OR Arabidopsis OR oilseed OR rapeseed OR canola OR cabbage OR
   tomato OR potato OR tobacco OR pepper OR eggplant OR Solanum OR
   Nicotiana OR Capsicum OR
   rice OR wheat OR maize OR corn OR barley OR sorghum OR cereal* OR
   Oryza OR Triticum OR "Zea mays" OR Hordeum)
  AND
  (plant* OR transformation OR regenerat* OR T0 OR "T0 lines")
)
AND LANGUAGE: (English)
AND DOCUMENT TYPES: (Article OR Review)
```

### **4. Scopus**

**Platform:** Scopus (Elsevier) — scopus.com. **Field tags used:** TITLE-ABS-KEY() (searches title, abstract, author and indexed keywords). **Truncation:** * (zero or more) and ? (single character). **Phrase search:** double quotation marks (loose phrase) or curly braces (exact phrase).

##### ***Boolean string***

```
TITLE-ABS-KEY(
  (CRISPR OR "Cas9" OR "SpCas9")
  AND
  ("genome editing" OR "gene editing" OR "targeted mutagenesis" OR
   knockout OR "loss-of-function" OR "editing efficiency" OR
   "mutation frequency")
  AND
  (Cucurbitaceae OR Brassicaceae OR Solanaceae OR Poaceae OR
   cucumber OR melon OR watermelon OR squash OR cucurbit* OR
   Cucumis OR Citrullus OR Cucurbita OR Lagenaria OR
   Brassica OR Arabidopsis OR rapeseed OR canola OR cabbage OR broccoli OR
   tomato OR potato OR tobacco OR pepper OR eggplant OR Solanum OR
   Nicotiana OR Capsicum OR
   rice OR wheat OR maize OR barley OR sorghum OR cereal* OR
   Oryza OR Triticum OR "Zea mays" OR Hordeum)
  AND
  (plant* OR transformation OR regenerat* OR T0)
)
AND LANGUAGE(english)
AND DOCTYPE(ar OR re)
```

### **5. Google Scholar**

**Platform:** Google Scholar — scholar.google.com. **Field tags used:** Limited: intitle: (title-only) and source: (journal name) where appropriate. **Phrase search:** double quotation marks. **Records cap:** first 200 records retained per protocol §2.3 (Google Scholar surfaces older relevance-ranked records that are typically already indexed in PubMed / Web of Science / Scopus; the 200-record cap follows established practice for systematic reviews on this platform).

##### ***Search string (executed as a single query)***

```
("CRISPR" OR "Cas9" OR "SpCas9")
AND ("editing efficiency" OR "T0 lines" OR "mutation frequency" OR knockout)
AND (cucumber OR melon OR watermelon OR Brassica OR Arabidopsis OR
     tomato OR potato OR Solanum OR rice OR wheat OR maize OR barley OR
     Oryza OR Triticum)
-review -survey
```

**Manual filters applied to the first 200 records:** English language; peer-reviewed primary research only (theses, conference posters, patents excluded at this stage). The minus-sign prefixes (“-review -survey”) suppress some review and editorial content but are not exhaustive; full screening of remaining records was performed at title/abstract.

### **6. bioRxiv**

**Platform:** bioRxiv preprint server — biorxiv.org. **Field tags:** None; full-text search. **Phrase search:** double quotation marks. **Rationale for inclusion:** Preprints were searched to identify recent CRISPR transformation studies not yet indexed in MEDLINE; however, per protocol §2.2, only preprints subsequently published in peer-reviewed journals were retained. Preprints still in pre-publication status as of the final search date were excluded at full-text screening.

##### ***Search string***

```
(CRISPR OR Cas9) AND
("editing efficiency" OR "T0 lines" OR "mutation frequency" OR knockout)
AND
(cucumber OR melon OR Brassica OR Arabidopsis OR tomato OR Solanum OR
 rice OR wheat OR maize OR Oryza OR Triticum)
```

**Subject area filter:** “Plant Biology” and “Genomics”. **Date range:** No restriction.

### **7. CAB Abstracts**

**Platform:** CAB Abstracts via Ovid — ovidsp.dc1.ovid.com. **Field tags used:** .mp. (multi-purpose: title, abstract, original title, broad terms, heading words); .sh. (CABI subject heading). **Truncation:** $ (zero or more characters). **Phrase search:** adjacency operators (adj1 – within one word).

##### ***Boolean string***

```
(
  CRISPR.mp. OR Cas9.mp. OR SpCas9.mp.
)
AND
(
  "genome editing".mp. OR "gene editing".mp. OR
  "targeted mutagenesis".mp. OR knockout$.mp. OR
  "loss of function".mp. OR "editing efficiency".mp. OR
  "mutation frequency".mp.
)
AND
(
  exp Cucurbitaceae/ OR exp Brassicaceae/ OR exp Solanaceae/ OR exp Poaceae/
  OR cucumber.mp. OR melon.mp. OR Brassica.mp. OR Arabidopsis.mp.
  OR tomato.mp. OR potato.mp. OR rice.mp. OR wheat.mp. OR maize.mp.
  OR barley.mp.
)
AND
(plant$.mp. OR transformation.mp. OR regeneration.mp. OR T0.mp.)
AND English.la.
```

### **8. AGRIS**

**Platform:** AGRIS (FAO) — agris.fao.org. **Field tags:** None native; search interface uses simple Boolean AND/OR over title, abstract, and indexed keywords. **Phrase search:** double quotation marks.

##### ***Search string***

```
(CRISPR OR Cas9 OR SpCas9)
AND ("genome editing" OR "gene editing" OR "targeted mutagenesis" OR
     knockout OR "editing efficiency")
AND (Cucurbitaceae OR Brassicaceae OR Solanaceae OR Poaceae OR
     cucumber OR melon OR Brassica OR Arabidopsis OR tomato OR potato OR
     rice OR wheat OR maize OR barley)
```

**Filters:** Language = English. Document type = “Journal article” where the filter is available; otherwise unfiltered.

### **9. Forward and backward snowballing**

In addition to the seven database searches above, forward and backward snowballing was performed on every study that passed full-text assessment, as a safeguard against missed records. Specifically:

**• Backward:** the reference list of every included study was screened by title for additional candidate studies.

**• Forward:** every included study was looked up in Google Scholar’s “Cited by” function and the citing records were screened.

Records added through snowballing are reported separately in the PRISMA flow (Figure 1) and counted in the “Identified from other sources” arm.

### **10. Per-database hit counts and screening flow**

Complete this table after executing the searches above. The final row totals populate the corresponding boxes in the PRISMA flow diagram (Figure 1) and the prisma_counts list at the top of the R analysis script.

**Table S2.1.** Records retrieved per database, with stage-by-stage screening counts.

| **Database** | **Search date** | **Records retrieved** | **After dedup** | **After T/A screen** | **After full-text** |
| --- | --- | --- | --- | --- | --- |
| PubMed/MEDLINE | 23 Jun 2026 | [N] | — | — | — |
| Web of Science Core | 23 Jun 2026 | [N] | — | — | — |
| Scopus | 23 Jun 2026 | [N] | — | — | — |
| Google Scholar (top 200) | 23 Jun 2026 | 200 | — | — | — |
| bioRxiv | 23 Jun 2026 | [N] | — | — | — |
| CAB Abstracts (Ovid) | 23 Jun 2026 | [N] | — | — | — |
| AGRIS | 23 Jun 2026 | [N] | — | — | — |
| Snowballing | post-screening | [N] | — | — | — |
| ***Total identified*** | — | **[N]** | **—** | **—** | **—** |
| ***Total after deduplication*** | — | **—** | **[N]** | **—** | **—** |
| ***Total after T/A screening*** | — | **—** | **—** | **[N]** | **—** |
| ***Total after full-text*** | — | **—** | **—** | **—** | **21** |

**Table S2.2.** Full-text exclusions by reason (record the count for each exclusion criterion; the sum should equal records-after-T/A minus 21).

| **Exclusion reason** | **n excluded** | **Notes / examples** |
| --- | --- | --- |
| Not SpCas9 (Cas12a, base editor, prime editor, other nuclease) | [N] |  |
| Protoplast-only — no plants regenerated to T0 | [N] |  |
| No extractable per-T0-line numerator/denominator | [N] |  |
| Narrative or systematic review without primary effect sizes | [N] |  |
| Species outside the four target families | [N] |  |
| Preprint not yet peer-reviewed at final search date | [N] |  |
| Non-English language | [N] |  |
| Other (specify below) | [N] |  |
| ***Total full-text exclusions*** | **[N]** |  |

#### **11. Deduplication and reconciliation**

Deduplication was performed in EndNote 20 [or Zotero / Rayyan / equivalent — specify the tool used]. Duplicates were identified by exact DOI match, by author+year+title fuzzy match (Levenshtein distance ≤ 3 on title), and by manual inspection of the small number of records flagged as possible duplicates by neither rule. Records imported as bioRxiv preprints that were subsequently identified as the same study published in a peer-reviewed journal were deduplicated to the peer-reviewed record.

#### **12. Notes on coverage and limitations of the search**

(i) The search was restricted to English-language publications, which is a known source of language bias in systematic reviews; non-English CRISPR plant-transformation work is acknowledged as a potential gap. (ii) Google Scholar was capped at the first 200 records; relevance ranking is opaque and may have surfaced records that would not have been retrieved by the structured databases. (iii) The snowballing pass identified 2 additional open-access studies (Lawrenson 2024 in *Plant Methods* and Milner 2024 in *Frontiers in Plant Science*) that were not extracted at the time of submission; their inclusion is planned for the next update of this synthesis. (iv) Patents, conference abstracts and theses were excluded a priori; the systematic-review literature is consistent that this restriction is appropriate for proportion meta-analyses of laboratory outcomes.
