## Supplementary material for "Editing Efficiency Across Crop Families: A Systematic Review and Meta-Analysis of CRISPR/SpCas9 Knockout Outcomes in Cucurbitaceae, Brassicaceae, Solanaceae and Poaceae": https://osf.io/vgf5u/files/dvm3g

**Supplementary File S5**

**Study-level details for all 23 included studies**

*Per-study summary of crop family, species, ploidy, delivery system, target class, detection assay, and contribution to primary versus sensitivity synthesis*

This file tabulates the 23 studies entering qualitative synthesis from the v5 PRISMA-2020 systematic review. Twenty-two studies contribute primary effect sizes (k = 172 total) and one (Ma X 2015) contributes per-locus rows reserved for sensitivity analysis. Three Zhang Z 2019 rows from the T1 generation are similarly sensitivity-only (see column *Sensitivity k*).

Family-level totals: Cucurbitaceae 14 effect sizes from 6 studies; Brassicaceae 20 from 4 studies; Solanaceae 68 from 4 studies (58 from Zhang N 2019); Poaceae 70 primary effect sizes from 8 studies (plus 1 sensitivity-only study, Ma X 2015). Full per-row metadata is in the extraction workbook *Data_Extraction_v5_EXTENDED.csv* deposited at OSF.

### **Per-study summary table**

| **ID** | **First author / year** | **Family** | **Species** | **Ploidy** | **Delivery** | **Primary k** | **Sensitivity k** | **Target class** | **Detection assay** |
| --- | --- | --- | --- | --- | --- | --- | --- | --- | --- |
| S001 | Tian 2017 | Cucurbitaceae | C. lanatus | Diploid | Agro | 1 | 0 | PDS | Sanger |
| S002 | Hooghvorst 2019 | Cucurbitaceae | C. melo | Diploid | Agro | 1 | 0 | PDS | Sanger |
| S003 | Pan 2022 | Cucurbitaceae | C. lanatus | Diploid | Agro | 1 | 0 | Quality | Sanger |
| S005 | Wang 2024 | Cucurbitaceae | C. melo | Diploid | Agro | 5 | 0 | Dev | Amplicon NGS |
| S006 | Xin 2022 | Cucurbitaceae | C. melo | Diploid | Agro | 3 | 0 | Dev | Amplicon NGS |
| S007 | Feng 2021 | Cucurbitaceae | C. lanatus | Diploid | Agro | 3 | 0 | Visible-marker | Sanger + NGS |
| S009 | Yang H 2017 | Brassicaceae | B. napus | Allotetraploid | Agro | 10 | 0 | Visible-marker | Sanger + T7E1 |
| S010 | Ma C 2019 | Brassicaceae | B. oleracea v. capitata | Allotetraploid | Agro | 4 | 0 | Visible-marker | Sanger |
| S013 | Yang Y 2018 | Brassicaceae | B. napus | Allotetraploid | Agro | 4 | 0 | Dev (CLAVATA) | PAGE + Sanger |
| S014 | Zheng 2020 | Brassicaceae | B. napus | Allotetraploid | Agro | 2 | 0 | Quality | Sanger |
| S015 | Brooks 2014 | Solanaceae | S. lycopersicum | Diploid | Agro | 3 | 0 | Dev (PROCERA) | Sanger + phenotype |
| S016 | Pan 2016 | Solanaceae | S. lycopersicum | Diploid | Agro | 4 | 0 | Multi-target | Sanger |
| S017 | Zhang N 2019 | Solanaceae | S. lycopersicum | Diploid | Agro | 58 | 0 | Disease resistance screen | Sanger + TIDE |
| S018 | Gonzalez 2021 | Solanaceae | S. tuberosum | Autotetraploid | Protoplast PEG/RNP | 3 | 0 | Quality (VInv) | HRFA + Sanger |
| S004 | Howells 2018 | Poaceae | T. aestivum | Hexaploid | Agro | 2 | 0 | Hormone (DEP1) | Sanger |
| S019 | Gasparis 2018 | Poaceae | H. vulgare | Diploid | Agro | 4 | 0 | Quality (naked grain) | PCR/RE + Sanger |
| S022 | Ma X 2015 | Poaceae | O. sativa | Diploid | Agro | 0 | 4 | Multi-target | Sanger + DSDecode |
| S023 | Zhang H 2014 | Poaceae | O. sativa | Diploid | Agro | 11 | 0 | Visible-marker (OsBADH2) | Sanger |
| S024 | Char 2017 | Poaceae | Z. mays | Diploid | Agro | 4 | 0 | Multi-target | T7E1 + Sanger |
| S025 | Lee 2019 | Poaceae | Z. mays | Diploid | Agro | 2 | 0 | Quality (waxy) | Sanger + TIDE |
| S026 | Zhang Z 2019 | Poaceae | T. aestivum | Hexaploid | Agro | 1 | 3 | Quality (gluten) | PCR-RE + Sanger |
| S027 | Lawrenson 2024 | Poaceae | H. vulgare + T. aestivum | Diploid + Hexaploid | Agro | 16 | 0 | Multi (Cas9+Cas12a toolkit) | PCR + Sanger |
| S028 | Milner 2024 | Poaceae | O. sativa + H. vulgare + T. aestivum | Diploid + Hexaploid | Agro | 30 | 0 | Hormone (GSK1) | PCR + Sanger |

**Notes.** Species abbreviations: *C. lanatus* = *Citrullus lanatus* (watermelon); *C. melo* = *Cucumis melo* (melon); *B. napus* = *Brassica napus* (oilseed rape); *B. oleracea v. capitata* = *Brassica oleracea* var. *capitata* (cabbage); *S. lycopersicum* = *Solanum lycopersicum* (tomato); *S. tuberosum* = *Solanum tuberosum* (potato); *H. vulgare* = *Hordeum vulgare* (barley); *T. aestivum* = *Triticum aestivum* (bread wheat); *O. sativa* = *Oryza sativa* (rice); *Z. mays* = *Zea mays* (maize). "Agro" = *Agrobacterium tumefaciens*-mediated stable transformation. "k" = effect sizes contributing to the named analysis (primary = pooled in main GLMM; sensitivity = reserved per pre-registered protocol). The Gonzalez 2021 row reports two effect sizes from PEG-mediated DNA delivery and one from RNP delivery; these are aggregated as "Protoplast PEG/RNP" in this summary. Lawrenson 2024 and Milner 2024 span multiple Poaceae species in a single study; the species column lists all member species and the ploidy column lists the corresponding ploidies.

### **Excluded studies log (40 full-text exclusions)**

Per-study reasons for the 40 full-text exclusions are tabulated by category in Figure 1 of the main manuscript: Not SpCas9 (n = 14), Protoplast-only with no plants regenerated (n = 8), No extractable per-T0-line numerator/denominator (n = 8), Review without primary data (n = 5), Species outside the four target families (n = 2), and Other (n = 3; 1 non-English, 2 not-yet-peer-reviewed preprints at the search date). A per-record exclusion log with PMID/DOI for each excluded study is deposited at the OSF project as *Excluded_Studies_with_Reasons.csv* and is not reproduced here for space reasons.
