## Supplementary material for "Editing Efficiency Across Crop Families: A Systematic Review and Meta-Analysis of CRISPR/SpCas9 Knockout Outcomes in Cucurbitaceae, Brassicaceae, Solanaceae and Poaceae": https://osf.io/vgf5u/files/v4tse

**Supplementary File S4**

**Moderator harmonisation rules**

*How raw values reported in the 23 included studies were mapped to the harmonised moderator levels used in the meta-analytic synthesis*

### **1. Purpose and scope**

The 23 included studies report transformation methodology in heterogeneous prose. To synthesise across studies, each moderator was reduced to a small number of harmonised categorical levels using the rules in this document. The rules were pre-specified before extraction began (see protocol, OSF deposit) with two minor refinements added post-hoc to handle the v5 extension studies (Lawrenson 2024 architecture-letter labelling; Milner 2024 guide-architecture contrast); both are flagged below.

Every effect size in the extraction workbook (*Data_Extraction_v5_EXTENDED.csv*) carries the raw value as reported by the source study (columns delivery, ploidy, detection_assay, sgRNA_promoter, target_class) and the harmonised category used for synthesis (the _clean variants used by the R analysis script). This file documents the mapping between the two.

### **2. Family (taxonomic family)**

Four levels, assigned from the species reported by each study and resolved against APG IV (2016). No reclassification was required for any of the 23 studies.

| **Harmonised level** | **Member species in this dataset** | **Studies (k)** |
| --- | --- | --- |
| Cucurbitaceae | Citrullus lanatus (watermelon), Cucumis melo (melon) | 6 |
| Brassicaceae | Brassica napus (oilseed rape), Brassica oleracea var. capitata (cabbage) | 4 |
| Solanaceae | Solanum lycopersicum (tomato), Solanum tuberosum (potato) | 4 |
| Poaceae | Triticum aestivum (wheat), Hordeum vulgare (barley), Oryza sativa (rice), Zea mays (maize) | 8 (primary) + 1 (Ma X 2015 sensitivity-only) |

### **3. delivery_clean (Cas9 + sgRNA delivery system)**

Three harmonised levels, collapsing the variation in agronomy/transformation prose into the operationally meaningful categories used by the literature. The category **Agrobacterium** includes all reports of *Agrobacterium tumefaciens*-mediated stable transformation of any tissue (cotyledon, hypocotyl, immature embryo, leaf disc), regardless of strain (LBA4404, EHA105, GV3101, AGL1). The two protoplast categories distinguish DNA-templated from protein-templated approaches.

| **Raw delivery as reported** | **Harmonised level (delivery_clean)** | **Effect sizes (k)** |
| --- | --- | --- |
| Agrobacterium (all strains, all tissues) | Agrobacterium | 170 |
| Protoplast PEG transfection of Cas9 plasmid | Protoplast PEG | 1 |
| Protoplast PEG transfection of Cas9 ribonucleoprotein | Protoplast RNP | 1 |

*Note.* Both protoplast effect sizes come from Gonzalez 2021 (potato). The non-significance of the univariate delivery Q-test (Q = 1.73, df = 2, p = 0.42) reflects the near-complete absence of independent variation in this column rather than equivalence of delivery systems. This is discussed in 3.3 and the Figure 4 caption of the main manuscript.

### **4. ploidy_clean (genome ploidy of the host species)**

Four harmonised levels, assigned from accepted cytogenetic descriptions of each species. Note that ploidy and family are confounded by construction in this dataset (hexaploid = Poaceae only; allotetraploid = Brassicaceae only); this is discussed in 4.2 of the main manuscript.

| **Species** | **Ploidy** | **Harmonised level (ploidy_clean)** |
| --- | --- | --- |
| Solanum lycopersicum, Cucumis melo, Citrullus lanatus, Oryza sativa, Hordeum vulgare, Zea mays | 2n=2x (diploid) | Diploid |
| Solanum tuberosum (potato) | 2n=4x (autotetraploid) | Autotetraploid |
| Brassica napus, Brassica oleracea var. capitata | 2n=4x (allotetraploid) | Allotetraploid |
| Triticum aestivum (wheat) | 2n=6x (allohexaploid; AABBDD) | Hexaploid |

### **5. detection_assay_clean (molecular detection method)**

Five harmonised levels. Where studies reported a hierarchical assay pipeline (e.g. PCR/RE pre-screen followed by Sanger sequencing for confirmation), the highest-resolution assay reported was used. Where multiple independent assays were combined (e.g. T7E1 plus Sanger), the assay generating the reported numerator was used.

| **Raw assay description (representative)** | **Harmonised level (detection_assay_clean)** | **Effect sizes (k)** |
| --- | --- | --- |
| Sanger sequencing alone; Sanger of PCR amplicons; Sanger with clone subset | Sanger | approx. 90 |
| PCR/RE (restriction enzyme cleavage) followed by Sanger | PCR-RE + Sanger | approx. 30 |
| T7 endonuclease I assay alone or as pre-screen + Sanger | T7E1 + Sanger | 8 |
| Hi-TOM amplicon NGS (Wang 2024, Xin 2022) | Amplicon NGS | 8 |
| TIDE / DSDecode chromatogram decomposition; HRFA fragment analysis | Sanger + decomposition | approx. 36 |

*Note.* The Q-between test for detection assay was non-significant (Q = 5.59, df = 4, p = 0.23). Counts above are rounded for the harmonised summary; exact counts per level are computed at run-time from the workbook by the R script and are not pre-tabulated here.

### **6. sgRNA_promoter_clean (sgRNA promoter system)**

Six harmonised levels distinguishing Pol III promoter source (Arabidopsis, rice, wheat, other species) and Pol II / tRNA / ribozyme architectures used by some recent studies for multiplexing.

| **Raw promoter as reported** | **Harmonised level (sgRNA_promoter_clean)** |
| --- | --- |
| AtU6-26, AtU6-29, AtU3b, AtU3d (Arabidopsis Pol III) | AtU6 / AtU3 (Pol III) |
| TaU6, TaU3 (wheat Pol III); HvU3 (barley Pol III) | Wheat/Barley U6 (Pol III) |
| OsU3, OsU6 (rice Pol III); PU6.1, PU6.2 | Rice U6/U3 (Pol III) |
| AtU6 with tRNA-flanked architecture (polycistronic processing) | tRNA-spaced Pol III |
| CmYLCV, OsUbiquitin (Pol II promoters with ribozyme or tRNA processing) | Pol II + processing |
| Not specified in source text | Not specified |

*Note: v5 architecture letters.* Lawrenson 2024 introduced an architecture-letter labelling scheme (A, B, C, D) corresponding to combinations of promoter source and processing strategy. These were mapped post-hoc to the levels above using the explicit descriptions in Lawrenson 2024 Table 1; the mapping is recorded in the extraction workbook (notes column) for every row. Milner 2024 contrasts Pol II + ribozyme against tRNA-spaced Pol III within the same gene targets; both arms are coded explicitly using the levels above. The univariate Q-between test for sgRNA_promoter_clean was highly significant (Q = 43.30, df = 5, p < 0.0001) but does not survive cluster-robust adjustment in the joint moderator model ( 3.4).

### **7. target_class_clean (functional class of the targeted gene)**

Five harmonised levels grouping target genes by their functional category. Assignment based on the gene description in the source study and TAIR / RAP-DB / Sol Genomics Network annotations.

| **Harmonised level** | **Examples of targeted genes** |
| --- | --- |
| Visible-marker / pigment | PDS (phytoene desaturase), CRTISO, ALB1; targets producing visible albino or carotenoid phenotype |
| Disease resistance / susceptibility | S genes (MLO, DMR6, SlORG); R genes; Zhang N 2019 disease-resistance screen |
| Hormone signalling / development | CLAVATA family (CLV1, CLV2, CLV3, CLE9), BRI1, GSK1, agriculturally relevant developmental QTLs |
| Quality / metabolite | GBSS (waxy starch), VInv (vacuolar invertase), homoeologue gluten genes |
| Other / reporter only | GFP, mCherry constructs without phenotypic target |

### **8. generation_clean (plant generation in which edits were scored)**

Two operational levels, with strict pre-registered handling of T1-generation data.

| **Raw generation as reported** | **Harmonised level** | **Pre-registered handling** |
| --- | --- | --- |
| T0 regenerants (first transformed plants) | T0 | PRIMARY synthesis |
| T1 progeny (segregating from T0) | T1 | SENSITIVITY only; not pooled with T0 rates |

*Note.* Per the pre-registered protocol, T1-generation edit rates are not interchangeable with T0 rates because T1 segregation can confound the editing-efficiency signal with Mendelian inheritance of the T0 edit. Three Zhang Z 2019 T1-wheat rows initially extracted as primary were reclassified to sensitivity during dual-reviewer reconciliation; this is documented in Supplementary File S8.

### **9. per_line_or_locus (outcome denominator type)**

Two strict operational levels with binary handling.

| **Raw outcome as reported** | **Harmonised level** | **Pre-registered handling** |
| --- | --- | --- |
| Number of T0 lines with confirmed edit / total T0 lines screened | per_line | PRIMARY outcome; pooled in the binomial-normal GLMM |
| Number of edited loci / total loci screened (per-locus rate) | per_locus | SENSITIVITY only; reserved for the sensitivity-with-locus analysis ( 3.5) |

*Note.* Four rows from Ma X 2015 (rice) are per-locus and excluded from primary synthesis. The primary_outcome_flag column in the workbook is the gating variable consumed by the R pipeline; rows with flag = “Yes” enter primary synthesis, rows with flag = “No” enter sensitivity only.

### **10. Missing data: handling and reconstruction**

Two operational rules applied during extraction:

**Zero numerators:** Studies reporting 0 edited T0 lines from *n* screened are extracted with numerator = 0 and the full denominator. No continuity correction is applied because the binomial-normal GLMM (rma.glmm with measure = PLO) handles zero counts exactly via the binomial likelihood. This is the principled handling of zero cells for proportion meta-analyses (Schwarzer et al. 2009, *Stat Med* 28:721–738).

**Percentage-only reporting:** Where a study reports only a percentage (e.g. “45% of T0 plants edited”) without raw counts, the numerator was reconstructed by multiplying the reported percentage by the reported denominator and rounding to the nearest integer. Two of the 46 v5 rows (Milner 2024 wheat tRNA guide 2, A and D homoeologues) have numerators reconstructed in this way; both are flagged in the workbook notes column. Reconstruction was applied only where the denominator was explicitly stated; where neither the numerator nor the denominator was extractable, the row was excluded at full-text stage (n = 8 of the 40 full-text exclusions; see Figure 1 of the main manuscript).

### **11. Reproducibility**

The harmonised levels in this document correspond directly to the _clean columns consumed by the R analysis script CRISPR_meta_analysis_v5.R (deposited at the OSF project). The mapping from raw text to harmonised level is encoded in the derive_clean_columns() function near the start of the script. To audit the mapping for any individual effect size, open the extraction workbook, locate the row by row_id, and compare the raw column (e.g. delivery) against the corresponding _clean derivation in the R function. Inter-rater agreement on every _clean column was κ ≥ 0.953 in the v4 dual-reviewer pass (Supplementary File S8); v5 verification of the 46 new rows is pending.
